# A *C. elegans* model of familial Alzheimer’s disease shows age-dependent synaptic degeneration independent of amyloid β-peptide

**DOI:** 10.1101/2025.07.16.665161

**Authors:** Vaishnavi Nagarajan, Caitlin L. Libowitz, Brian D. Ackley, Michael S. Wolfe

## Abstract

The membrane-embedded γ-secretase complex is involved in the intramembrane cleavage of ∼ 150 substrates. Cleavage of amyloid precursor protein (APP)-derived substrate C99 generates 38-43-residue secreted amyloid β-peptides (Aβ), with the aggregation-prone 42-residue form (Aβ42) particularly implicated in the pathogenesis of Alzheimer’s Disease (AD). However, whether Aβ42 is the primary driver of neurodegeneration in AD remains unclear. Dominant mutations in APP or presenilin—the catalytic component of γ-secretase—cause early-onset familial AD (FAD) and reduce one or more steps in the multi-step processive proteolysis of C99 to Aβ peptides, apparently through stabilization of γ-secretase enzyme-substrate (E-S) complexes. To investigate mechanisms of neurodegeneration in FAD, we developed new *C. elegans* models co-expressing wild-type or FAD-mutant C99 substrate and presenilin-1 (PSEN1) variants in neurons, allowing intramembrane processing of C99 to Aβ *in vivo.* We demonstrate that while FAD-mutation of either C99 or PSEN1 leads to age-dependent synaptic loss, proteolytically inactive PSEN1 did not. Designed mutations that allow stable E-S complex formation without Aβ42 or Aβ production likewise result in synaptic degeneration. Moreover, replacement of C99 with variants of a Notch1-based substrate revealed that disrupted processing of another γ-secretase substrate can similarly lead to synaptic degeneration. These results support a model in which synaptic loss can be triggered by toxic, stalled γ-secretase E-S complexes in the absence of Aβ production and not by simple loss of proteolytic function. This new *C. elegans* system provides a powerful platform to study the role of dysfunctional γ-secretase substrate processing in FAD pathogenesis.

**SIGNIFICANCE:** Dominantly inherited familial Alzheimer’s disease (FAD) is caused by mutations in the enzyme or substrate that produces amyloid β-peptides (Aβ). These mutations alter enzyme processing of substrate to Aβ and can skew production of Aβ to aggregation-prone forms that deposit as pathological plaques. Nevertheless, whether Aβ is the primary driver of AD remains unresolved. Recent evidence supports a model in which loss of neuronal connections in FAD is due to the stalled enzymatic process of Aβ production, rather than the Aβ products. Using the roundworm *C. elegans* as a genetic model system, we show here that the stalled process itself, rather than Aβ or reduced enzyme activity, can trigger loss of neuronal connections with age in this simple model animal.

## INTRODUCTION

Alzheimer’s disease is a progressive neurodegenerative disorder causing cognitive decline and dementia in over 50 million worldwide (Breijyeh and Karaman 2020). AD is pathologically characterized by cerebral deposition of two proteins: amyloid β-peptide (Aβ), forming extra-neuronal plaques, and the microtubule-associated protein tau, forming intraneuronal fibrillary tangles (Cai and Tammineni 2017). Synaptic damage and degeneration occur in early stages, eventually leading to neuronal loss (Pelucchi, Gardoni et al. 2022). Alterations in axonal transport of synaptic vesicles, mitochondrial damage, neuroinflammation, and oxidative stress leading to axonal dystrophy have been implicated in synaptic loss in AD (Krstic and Knuesel 2013, Subramanian, Savage et al. 2020).

Aβ aggregation has long been considered the initiator of synaptic damage and neurodegeneration, per the amyloid hypothesis of AD pathogenesis. Aβ is produced from APP by sequential proteolysis by β-secretase, which sheds the large lumenal/extracellular ectodomain, and γ-secretase, which cleaves the remnant 99-residue fragment (C99) within its single transmembrane domain (Zhang, Ma et al. 2012). Rare dominant missense mutations in APP and presenilin, the catalytic component of γ-secretase, cause early-onset familial AD (FAD)(Goate, Chartier-Harlin et al. 1991, Sherrington, Rogaev et al. 1995), with the same pathology, presentation and progression as the more common sporadic late-onset AD (Bateman, Aisen et al. 2011, Morris, Weiner et al. 2022), strongly suggesting common pathogenic mechanisms. These mutations alter Aβ production and properties, generally increasing the proportion of the aggregation-prone 42-residue peptide (Aβ42) over the more soluble 40-residue variant (Aβ40) (Tanzi 2012). Nevertheless, after over 30 years of the amyloid hypothesis, mechanisms by which Aβ aggregates trigger synaptic and neuronal degeneration remain unresolved (Makin 2018). Moreover, therapeutic candidates targeting Aβ repeatedly failed in clinical trials until the recent approval of monoclonal antibodies. These new antibodies, however, are controversial, due to concerns about efficacy and safety (Hunter 2024).

The lack of mechanistic understanding and limited effectiveness of therapeutics suggests that Aβ may not be the primary disease driver in AD. FAD mutations, however, are only found in genes encoding the substrate (APP) or enzyme (γ-secretase) that produce Aβ, pointing to alteration of proteolytic processing by γ-secretase in pathogenesis. Proteolysis of C99 by γ-secretase is complex: initial endoproteolysis produces long Aβ peptide intermediates Aβ48 or Aβ49 that are then processively trimmed, generally in tripeptide increments, to shorter secreted forms (Takami, Nagashima et al. 2009). We found that FAD mutations in APP and PSEN1 are consistently deficient in early processing steps, rather than later steps that produce Aβ40 and Aβ42 (Devkota, Williams et al. 2021, Devkota, Zhou et al. 2024, Arafi, Devkota et al. 2025, Devkota, Maesako et al. 2025). Moreover, we found that FAD mutations stabilize the enzyme-substrate complex, suggesting a mechanism for the reduced proteolytic function (Levitan, Doyle et al. 1996, Devkota, Zhou et al. 2024, Arafi, Devkota et al. 2025, Devkota, Maesako et al. 2025).

To test mechanisms by which FAD mutations trigger neurodegeneration, we developed a *C. elegans* transgenic model system that expresses C99 and/or PSEN1 in neurons (Devkota, Zhou et al. 2024). C99 includes a signal sequence for membrane insertion, and PSEN1 replaces *C. elegans* orthologs *sel-12* and *hop-1* to provide functional γ-secretase with the human catalytic component (Levitan, Doyle et al. 1996). Thus, in this system Aβ is produced from substrate and enzyme via intramembrane proteolysis, as it is normally, in contrast to previous *C. elegans* models that expressed Aβ directly and in the cytosol. Using this system, we found that FAD mutations lead to age-dependent synaptic degeneration and reduced lifespan. Moreover, this phenotype did not require Aβ production and implicated the stalled enzyme-substrate complex as the pathogenic trigger. Here we reproduce these findings in independent transgenic lines and extend these studies by showing that the neurodegenerative phenotype is not due to simple loss of proteolytic function of stalled complexes. We also test that ability of another γ-secretase substrate, Notch1, to induce the neurodegenerative phenotype. Taken together, these results suggest that stalled γ-secretase enzyme-substrate complexes *per se* trigger a novel gain of synaptotoxic function.

## RESULTS

Previous attempts to develop a *C. elegans* model for Alzheimer’s disease involved expression of the amyloid β-peptide (Aβ), particularly Aβ42, which pathologically deposits in the Alzheimer brain (Link 1995, Bieschke, Cohen et al. 2009, McColl, Roberts et al. 2009, Treusch, Hamamichi et al. 2011, McColl, Roberts et al. 2012). However, Aβ is normally produced in cellular membranes through sequential proteolysis of the amyloid precursor protein (APP), first by β-secretase to produce the membrane-bound C-terminal APP stub C99, followed by intramembrane cleavage by γ-secretase. Dominant FAD-causing missense mutations in the substrate or enzyme alter the proteolytic processing of APP by γ-secretase. An *in vivo* model that recapitulates the proteolytic processing of APP substrate, as it occurs naturally, is critical to understanding how FAD mutations contribute to pathogenesis. Such *C. elegans* models, in which Aβ production occurs through γ-secretase in membranes have not been previously described. For these reasons, we engineered novel *C. elegans* models that co-express membrane-associated human C99 and human PSEN1 in neurons to allow for intramembrane proteolysis to produce Aβ.

We designed transgenes that express human wt C99APP or human wt PSEN1 with the *rgef-1* promoter, which drives stable transgene expression with age pan-neuronally in *C. elegans* (Chen, Fu et al. 2011, Adamla and Ignatova 2015, Munoz-Jimenez, Ayuso et al. 2017). The APP N-terminal signal peptide sequence precedes the C99APP sequence to allow membrane insertion. *unc-54* 3’ UTR was used to promote transgenic expression in somatic cells (Merritt, Rasoloson et al. 2008). Additionally, C99APP and PSEN1 variant transgenes were created using the same expression vector (**Figure 1a**). Transgenic worms were obtained by microinjection of human C99APP and human PSEN1 constructs in the *juIs1* line (Hallam and Jin 1998), which expresses GFP-fused synaptobrevin protein in presynaptic terminals of GABAergic motor neurons, visualized as synaptic puncta along the dorsal and ventral nerve cords. Similarly, transgene encoding L166P FAD PS-1 mutation was designed to express pan-neuronally in *C. elegans*. Schematic representation γ-secretase tripeptide cleavage driven by three active site pockets (S1’, S2’ and S3’) of presenilin (PSEN) is shown along with the corresponding three amino acids (P1’, P2’ and P3’) of substrate in **Figure 1b**. We previously showed that co-expression of mutant C99APP and/or PSEN1 led to neurodegeneration phenotypes (Devkota, Zhou et al. 2024).

**Figure 1.**
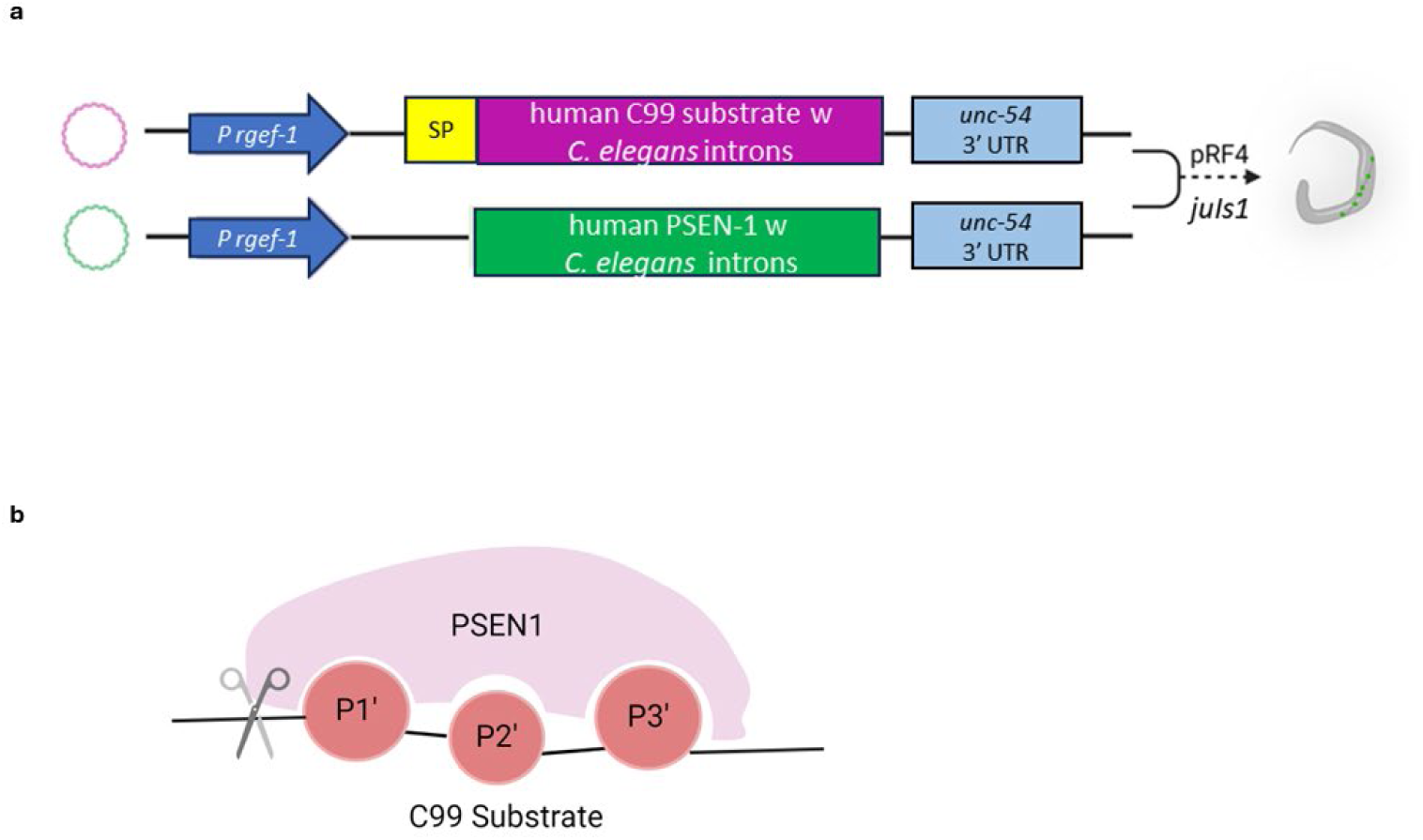
Design and creation of transgenic *C. elegans*. **a. Schematic of transgenic *C. elegans* vectors.** Transgenic *C. elegans* lines were obtained by microinjection of either of or combination of human PSEN-1(also called PS-1) and human C99 substrate (APP or Notch-1) driven by pan-neuronal *rgef-1* promoter into the parental line *juIs1. rol-6* (dominant roller marker) expressed in pRF-4 plasmid was used as the co-injection marker to select for transformants. This results in transformant *C. elegans* that are roller worms having synaptic puncta marked by GFP-tagged synaptobrevin along the nerve cord. **b. Visual representation of substrate processing by PSEN-1.** PSEN-1 proteolyzes substrates by tripeptide trimming. P1’, P2’ and P3’ are amino acids of the substrate accommodated in the 3 pockets of PSEN-1.

*C. elegans* lines co-expressing wt C99APP and wt PSEN1 showed lifespans that were comparable to those of the parental line *juIs1* **<u>(Figure S1a)</u>**. The log rank test of the survival curves showed that there is no statistically significant difference (P = 0.851). Numbers of synaptic puncta in transgenic lines were similar to those of the parental line for each day examined (Days 1-7 & 9) **(Figure S1b)**.

### wt PS-1 exacerbates I45F C99-induced synaptic loss in *C. elegans*

The Iberian mutation in APP, I45F (Aβ/C99 numbering), is an FAD mutation that places Phe in the P2’ position relative to Aβ46→Aβ43 tripeptide trimming. We have previously demonstrated that Phe in the P2’ position with respect to any cleavage event in the processive proteolysis of C99 substrate by γ-secretase blocks that cleavage step. Thus, for I45F the production of Aβ43 is prevented while allowing Aβ45→Aβ42 cleavage, thereby increasing the Aβ42/Aβ40 ratio (Bolduc, Montagna et al. 2016) (**Figure 2a**). The median lifespan of both of I45F C99APP-expressing lines (with and without coexpression of wt PSEN1) were shorter than wt C99APP + wt PSEN1 animals, although wt PSEN1 coexpression resulted in substantial reduction in lifespan and early loss of synapses (**Figure 2b; Table S1)**. *C. elegans* lines that express I45F C99APP and wt PSEN1 showed loss of synaptic puncta beginning on day 3 in dorsal and ventral nerve cords, with substantial synaptic degeneration occurring by day 9 in the dorsal cord. In contrast, the monogenic line expressing I45F C99APP alone showed only slight loss of synaptic puncta in the dorsal cord on day 9. (**Figure 2c-d**). We observed similar results in additional independent lines with the same transgenes, as reported in our previous study (Devkota, Zhou et al. 2024). These results indicate that I45F C99APP induces a weak neurodegenerative phenotype—likely due to interaction with endogenous presenilin sel-12 and/or hop-1—that is strongly exacerbated by coexpression of wt human PS-1, demonstrating a genetic interaction between the two transgenes.

**Figure 2.**
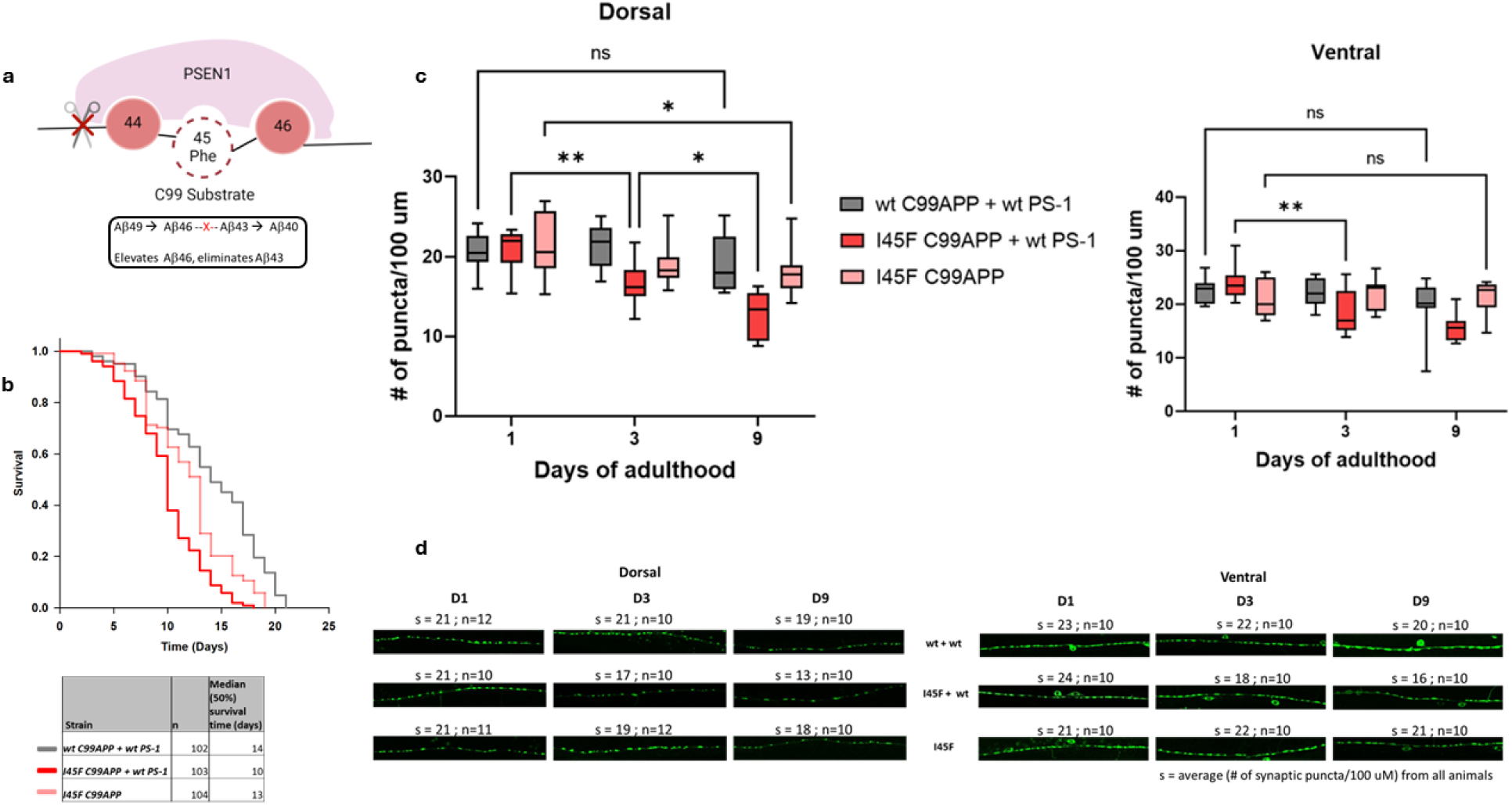
wt PS-1 exacerbates I45F C99-induced synaptic loss in *C. elegans*. **a. Visual representation of I45F C99APP processing by PSEN-1.** C99APP has a Phe substitution in the P2’ position of the cleavage site preventing the formation of Aβ43. **b. Shorter lifespan of I45F C99APP + wt PS-1 animals.** Kaplan-Meier curves of ‘n’ animals show the fraction of animals alive on different days. Ages of control animals - wt C99APP + wt PS-1 and I45F C99APP at median survival were 14 & 13 days respectively, while that of I45F C99APP + wt PS-1 was significantly shorter (10 days). **c. Age-dependent loss of synaptic puncta in I45F C99APP + wt PS-1 animals.** The average number of synaptic puncta per 100 μm in ‘n’ transgenic worms for each day is shown in the vertical box plots for dorsal and ventral cords. The horizontal lines in the top and bottom of any box plot represent the minimum and the values in that dataset of ‘n’ values. The ‘n’ for each day for a given mutant can be found in panel d. The upper and lower ends of a box mark the quartiles Q1 and Q3 values respectively. The horizontal line inside the box shows the median value (also, 2^nd^ quartile Q2). The number of synaptic puncta in I45F C99APP + wt PS-1 reduced on day 3 compared to day 1 significantly in dorsal nerve cord, while loss of synaptic puncta in I45F C99APP animals was first seen later on day 9. (p>0.05 ns, p≤ 0.05 *, p≤ 0.01 **, p≤ 0.001 ***, p< 0.0001 ****). Similarly, in ventral nerve cord, synaptic density in I45F C99APP + wt PS-1 reduced significantly on day 3 compared to day 1 but no significant reduction in the same was seen in either control. d. **Age-dependent loss of synaptic puncta in I45F C99APP + wt PS-1 animals.** 100 μm sections of representative confocal images for wildtype (1^st^ row) and transgenic animals are shown for days 1, 3 and 9 for dorsal (left) and ventral (right). ‘s’ denotes the average of (number of synaptic puncta per 100 μm) from ‘n’ animals. In I45 C99APP + wt PS-1 dorsal nerve cord, on day 3, s value dropped to17, compared to s=21 on day 1. Similarly, the value of s dropped to 18 on day 3 compared to 24 on day 1 in ventral nerve. Note the gaps in synaptic puncta in these images, increasing the distance between two consecutive puncta.

### Synaptic loss is triggered independently of Aβ42

Addition of a designed V44F mutation to the Iberian FAD-mutant C99APP (V44F/I45F C99APP) eliminates the production of Aβ42 by blocking processive proteolysis by γ-secretase at the Aβ45→Aβ42 step, as Phe is located in the P2’ position relative to this cleavage site, along with blocking Aβ46→Aβ43 due to I45F (**Figure 3a**) (Pope, Wilkins et al. 2021, Devkota, Zhou et al. 2024). Thus, this double mutant can address whether Aβ42 is required for the synaptic degeneration observed with I45F mutation alone. The median lifespan of V44F/I45F C99 APP + wt PSEN1 was substantially shorter than wt C99APP + wt PSEN1, while expression of V44F/I45F alone attenuated this effect (**Figure 3b**). The survival curves of wt C99APP + wt PS-1 and V44F/I45F C99 APP + wt PSEN1 were significantly different based on Holm-Sidak pairwise comparison **(Table S1)**. Similarly, significant loss of synaptic puncta was seen in dorsal and ventral nerve cords on day 2 in lines neuronally co-expressing V44F/I45F C99APP and wt PSEN1. A less severe phenotype with a delayed onset and absence of further synaptic loss was seen in the monogenic V44FI45F C99APP line (i.e., age-dependent loss of synaptic puncta was not observed in this control) (**Figure 3c-d**), recapitulating observations of an independent line with same mutations (Devkota, Zhou et al. 2024). These observations collectively suggest that Aβ42 is not essential for the neurodegenerative phenotype of these lines.

**Figure 3.**
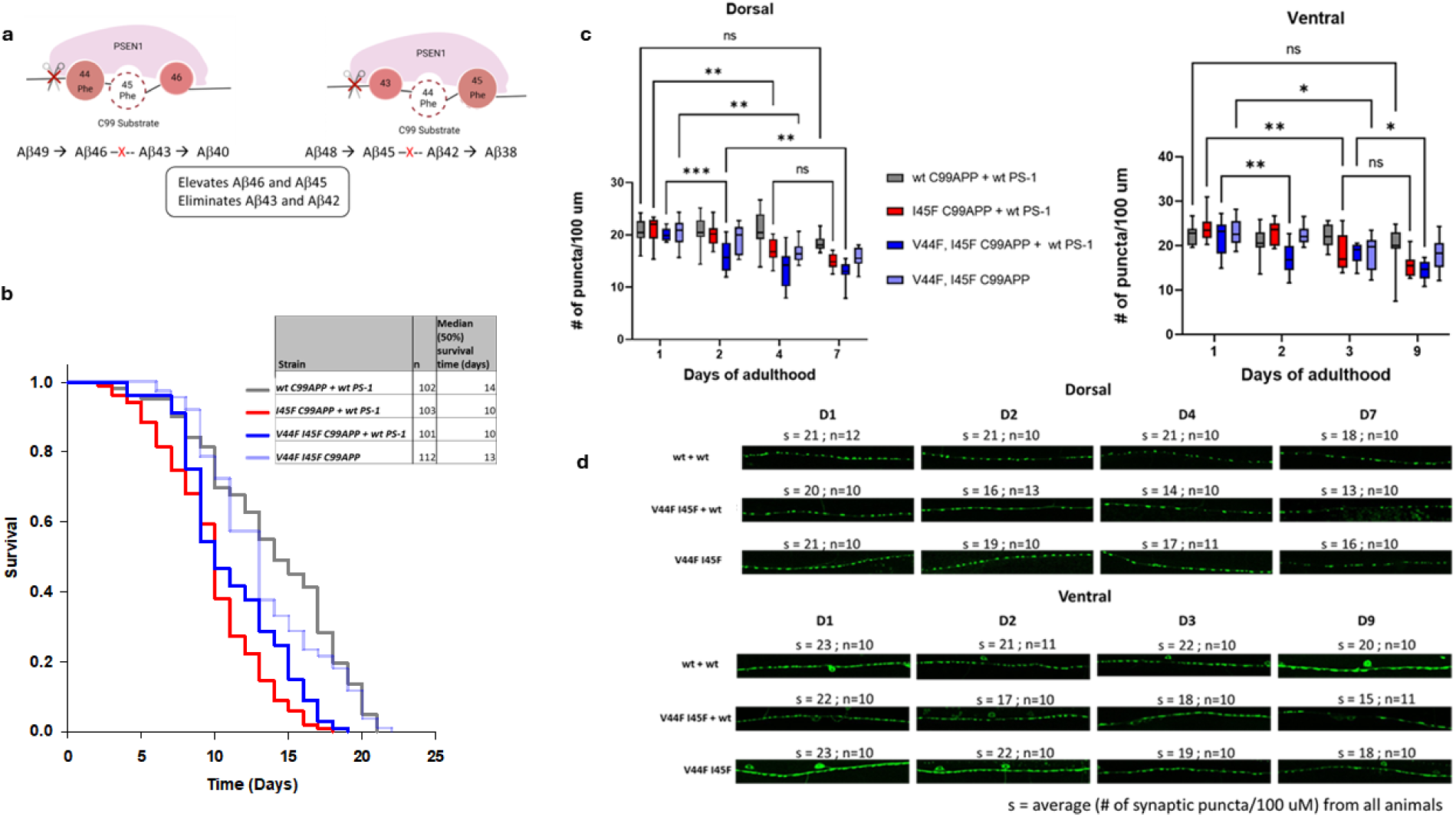
Synaptic loss is triggered independently of Aβ42. **a. Visual representation of V44F I45F C99APP processing by PSEN-1.** C99APP has two Phe substitutions in the P2’ positions each along Aβ40 and Aβ42 pathways preventing the formation of Aβ43 and Aβ42. **b. Shorter lifespan of V44F I45F C99APP + wt PS-1 animals.** Kaplan-Meier curves of ‘n’ animals show the fraction of animals alive on different days. The ages of control animals (wt C99APP + wt PS-1 and V44F I45F C99APP) at median survival were 14 and 13 days respectively, while that of V44F I45F C99APP + wt PS-1 was significantly shorter (10 days). **c. Age-dependent loss of synaptic puncta in V44F I45F C99APP + wt PS-1 animals.** The average number of synaptic puncta per 100 μm in ‘n’ transgenic worms for each day is shown in the vertical box plots for dorsal and ventral cords. The horizontal lines in the top and bottom of any box plot represent the minimum and the values in that dataset of ‘n’ values. The ‘n’ for each day for a given mutant can be found in panel d. The upper and lower ends of a box mark the quartiles Q1 and Q3 values respectively. The horizontal line inside the box shows the median value (also, 2^nd^ quartile Q2). The number of synaptic puncta in V44F I45F C99APP + wt PS-1 reduced on day 2 compared to day 1 significantly in dorsal nerve cord and then again on day 7, while significant loss of synaptic puncta in control V44F I45F C99APP first occurred later on day 4. Loss of synaptic puncta in V44F I45F C99APP + wt PS-1 first occurred on day 3 compared to day 1, while that was first seen later on day 3 in V44F I45F C99APP animals’ ventral nerve. In control and in FAD mutant I45F C99APP + wt PS-1, loss of synaptic puncta occurred later in dorsal and ventral nerves than that in V44F I45F C99APP + PS-1 mutant. (p>0.05 ns, p≤ 0.05 *, p≤ 0.01 **, p≤ 0.001 ***, p< 0.0001 ****). **d. Age-dependent loss of synaptic puncta in V44F I45F C99APP + wt PS-1 animals.** 100 μm sections of representative confocal images for wildtype (1^st^ row) and transgenic animals are shown for days 1, 2, 4 and 7 for dorsal (left) and days 1, 2, 3 and 9 for ventral (right). ‘s’ denotes the average of (number of synaptic puncta per 100 μm) from ‘n’ animals. In V44F I45 C99APP + wt PS-1 dorsal nerve cord, on day 2, s value dropped significantly to16, compared to s=20 on day 1. Similarly, the value of s dropped significantly to 17 on day 2 compared to 22 on day 1 in ventral nerve. Note the gaps in synaptic puncta in these images, increasing the distance between two consecutive puncta.

### Synaptic loss is triggered independently of Aβ

V50F/M51F mutation in C99APP blocks the initial endoproteolytic (ε) cleavage that releases APP intracellular domain and generates intermediates A*β*_48_ and A*β*_49_, due to the presence of Phe in the P2’ pocket relative to either cleavage site. This prevents C99APP substrate from being processed to Aβ, thereby providing a substrate suitable for testing if synaptic degeneration is dependent on any form of Aβ (**Figure 4a**). While V50F/M51F C99APP not proteolyzed by γ-secretase, binding of this substrate to the protease is not affected (Bolduc, Montagna et al. 2016). Lifespan of animals expressing V50F/M51F C99APP + wt PSEN1 was reduced similarly to that of those coexpressing wt PSEN1 pIus either I45F C99APP or V44F/I45F C99APP, as measured by median survival times (**Figure 4b**). Pairwise comparison of the survival curves showed that there is a significant difference between wt C99APP + wt PSEN1 and V50FM51F C99APP + wt PSEN1 **(Table S1)**. A reduced number of synaptic puncta was seen in the dorsal cord of these lines from as early as day 1 of adulthood compared to wt + wt transgenic lines (**Figure 4c-d**). In contrast, co-expression of wt PSEN1 with V50FM51F C99APP led to fewer synaptic puncta in the ventral compared to wt control animals on day 3. However, an increase in synaptic puncta was observed in V50FM51F C99APP mutants from day 7 in both nerve cords, a phenomenon not seen in lines co-expressing I45F or V44FI45F C99APP with wt PS-1. This aligns with our observation of independent lines described in our previous study and confirms that prevention of Aβ formation does not rescue loss of synaptic puncta, while supporting mechanisms of regeneration in this model (Devkota, Zhou et al. 2024)

**Figure 4.**
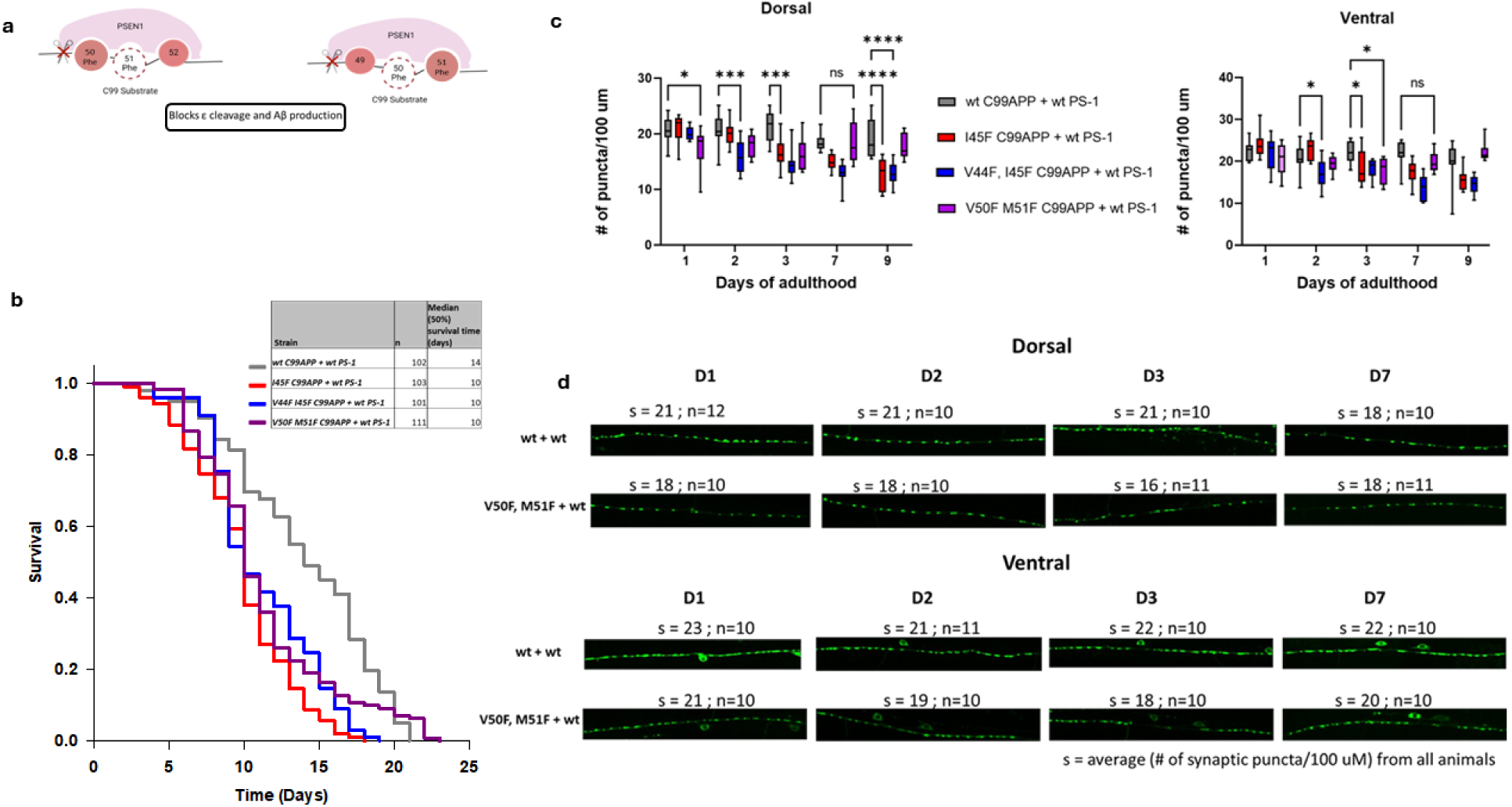
Synaptic loss is triggered independently of Aβ. **a. Visual representation of V50F M51F C99APP processing by PSEN-1.** C99APP has two Phe substitutions in the P2’ positions each along Aβ40 and Aβ42 pathways preventing the formation of Aβ production. **b. Shorter lifespan of V50F M51F C99APP + wt PS-1 animals.** Kaplan-Meier curves of ‘n’ animals show the fraction of animals alive at different days. Age of control animals wt C99APP + wt PS-1 at median survival was 14 days, while that of V50F M51F C99APP + wt PS-1 was significantly shorter (10 days). **c. Loss of synaptic puncta in V50F M51F C99APP + wt PS-1 animals.** The average number of synaptic puncta per 100 μm in ‘n’ transgenic worms for each day is shown in the vertical box plots for dorsal and ventral cords. The horizontal lines in the top and bottom of any box plot represent the minimum and the values in that dataset of ‘n’ values. The ‘n’ for each day for a given mutant can be found in panel d. The upper and lower ends of a box mark the quartiles Q1 and Q3 values respectively. The horizontal line inside the box shows the median value (also, 2^nd^ quartile Q2). The number of synaptic puncta in dorsal nerve cord of V50F M51F C99APP + wt PS-1 was significantly lower on day 1 (s= 18) compared to wt C99APP +wt PS-1 (s=21). Similar observations were seen until day 7 when there was no significant difference in s values between control and V50F M51F C99APP + wt PS-1. In case of ventral, on day 2 s=19 in V50F M51F C99APP + wt PS-1 was significantly less compared to s=21 in wt C99APP + wt PS-1. I45F C99APP + wt PS-1 and V44F I45F C99APP + wt PS-1 animals had significantly less s values compared to wt C99APP + wt PS-1 animals later on day 3 and day 2 respectively. On day 9, there was no significant difference in s values between wt C99APP + wt PS-1 and V50F M51F C99APP + wt PS-1 mutant. (p>0.05 ns, p≤ 0.05 *, p≤ 0.01 **, p≤ 0.001 ***, p< 0.0001 ****) **d. Loss of synaptic puncta in V50F M51F C99APP + wt PS-1 animals.** 100 μm sections of representative confocal images for wildtype (1^st^ row) and transgenic animals are shown for days 1, 2, 3 and 7 for dorsal (left) and ventral (right). ‘s’ denotes the average of (number of synaptic puncta per 100 μm) from ‘n’ animals. Note the gaps in synaptic puncta in these images, increasing the distance between two consecutive puncta.

### L166P PSEN1 induces synaptic loss independent of C99APP coexpression

The median lifespan in our new *C. elegans* lines expressing PSEN1 L166P FAD mutation, irrespective of the presence of C99APP substrate, was reduced compared to that of wt C99APP + wt PSEN1 animals (**Figure 5a; Table S1)**. wt C99APP + L166P lines also displayed age-dependent decline in the numbers of synaptic puncta in both cords (**Figure 5b,c**), results consistent with independent lines studied earlier (Devkota, Zhou et al. 2024). In our previous study, however, imaging of synaptic puncta in monogenic PSEN1 L166P lines was not possible due to toxic sensitivity to the anesthetic. This problem was solved here by immobilizing animals with polystyrene beads, which revealed that monogenic PSEN1 L166P *C. elegans* did not show age-dependent synaptic degeneration, with results indistinguishable from wt C99APP + wt PSEN1 lines (**Figure 6a,b**). These observations indicate that co-expression of exogenous substrate is necessary for synaptic degeneration, while the shortened lifespan is independent of substrate co-expression. This is the sole instance where shortened lifespan and age-dependent synaptic loss did not correlate, suggesting that, at least in the monogenic L166P PSEN1 lines, the two phenotypes are caused by distinct mechanisms.

**Figure 5.**
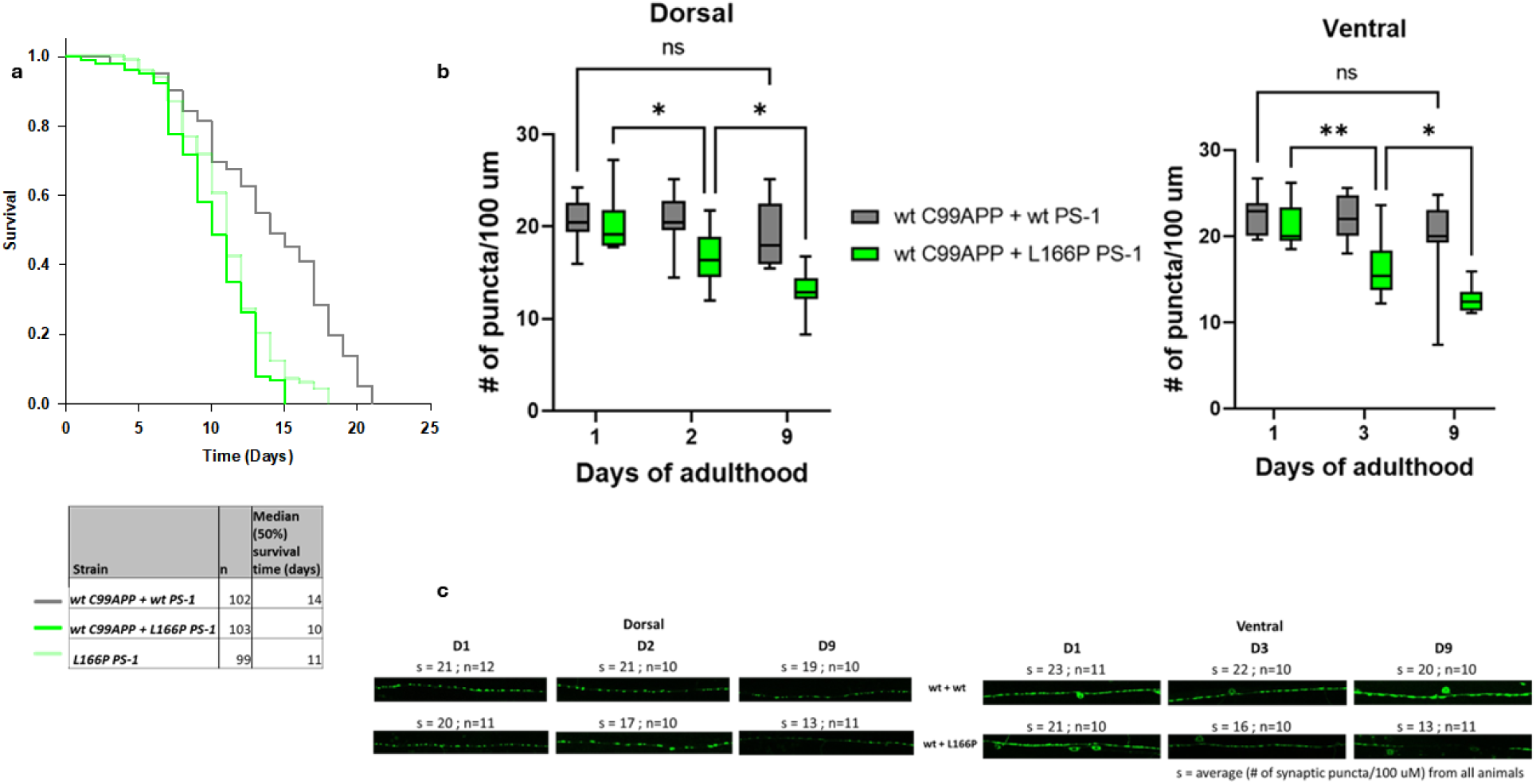
L166P PS-1 reduces lifespan independent of C99APP coexpression. **a. Shorter lifespan of wt C99APP + L166P PS-1 animals.** Kaplan-Meier curves of ‘n’ animals show the fraction of animals alive on different days. Age of control animals (wt C99APP + wt PS-1 at median survival was 14 days while that of wt C99APP + L166P PS-1 was significantly shorter (10 days). The median survival age of L166P PS-1 animals was 11 days. **b. Age-dependent loss of synaptic puncta in wt C99APP + L166P PS-1 FAD animals.** The average number of synaptic puncta per 100 μm in ‘n’ transgenic worms for each day is shown in the vertical box plots for dorsal and ventral cords. The horizontal lines in the top and bottom of any box plot represent the minimum and the values in that dataset of ‘n’ values. The ‘n’ for each day for a given mutant can be found in panel c. The upper and lower ends of a box mark the quartiles Q1 and Q3 values respectively. The horizontal line inside the box shows the median value (also, 2^nd^ quartile Q2). In comparison to day 1, the number of synaptic puncta in wt C99APP + L166P PS-1 FAD animals reduced on day 2 significantly in dorsal and day 3 in ventral nerve cords. Age dependent loss of synaptic puncta was seen in wt C99APP + L166P PS-1 FAD animals as the number dropped significantly compared to day 2 on day 9 in dorsal and ventral nerve cords. (p>0.05 ns, p≤ 0.05 *, p≤ 0.01 **, p≤ 0.001 ***, p< 0.0001 ****). **c. Age-dependent loss of synaptic puncta in wt C99APP + L166P PS-1 FAD animals.** 100 μm sections of representative confocal images for wildtype (1^st^ row) and transgenic animals are shown for days 1, 2 and 9 for dorsal (left) and 1, 3 and 9 for ventral (right). ‘s’ denotes the average of (number of synaptic puncta per 100 μm) from ‘n’ animals. In wt C99APP + L166P PS-1 dorsal nerve cord, on day 2, s value dropped significantly to17, compared to s=20 on day 1. Similarly, the value of s dropped significantly to 16 on day 3 compared to 21 on day 1 in ventral nerve. Note the gaps in synaptic puncta in these images, increasing the distance between two consecutive puncta.

**Figure 6.**
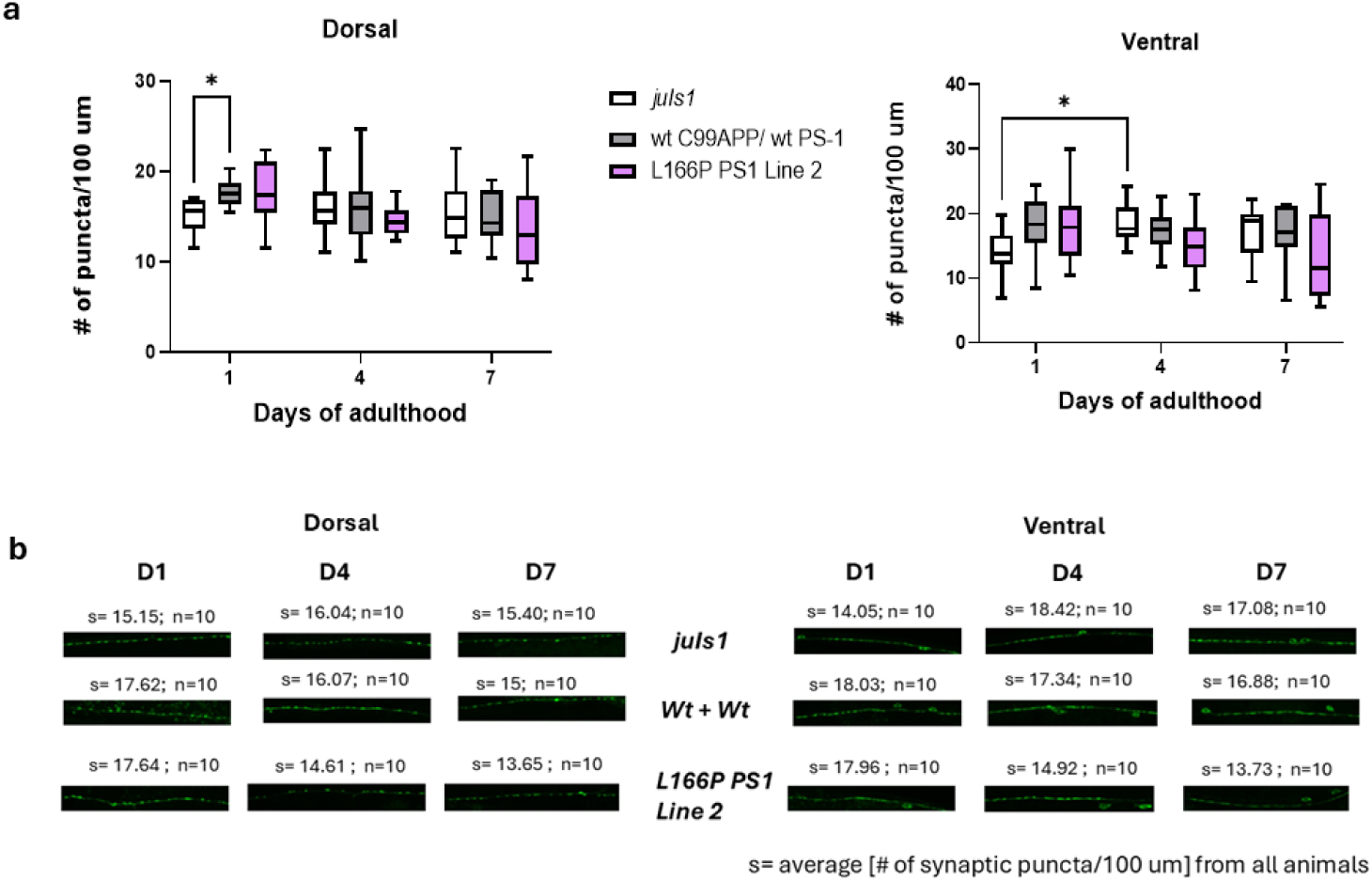
L166P without co-expression of WTC99 APP does not result in synaptic degeneration. **a. Synaptic puncta quantification in L166P PS-1 monogenic animals.** The average number of synaptic puncta per 100 μm in ‘n’ transgenic worms for each day is shown in the vertical box plots for dorsal and ventral cords. The horizontal lines in the top and bottom of any box plot represent the minimum and the values in that dataset of ‘n’ values. The ‘n’ for each day for a given mutant can be found in panel b. The upper and lower ends of a box mark the quartiles Q1 and Q3 values respectively. The horizontal line inside the box shows the median value (also, 2^nd^ quartile Q2). In comparison to wt C99APP + wt PS-1 control, the number of synaptic puncta in L166P monogenic animals is not significantly different on the days quantified (days 1, 4 & 7) in dorsal and ventral nerve cords. (p>0.05 ns, p≤ 0.05 *). There was some significant difference between controls: on day 1 of adulthood, *juIs1* was significantly decreased as compared to wt C99 APP + wt PS1-1 control (p=0.0121) for dorsal, and significantly different between days 1 and 4 within the strain (p=0.0216) for ventral. **b. Synaptic puncta quantification of L166P PS-1 monogenic animals.** 100 μm sections of representative confocal images for controls (*juIs1* - 1^st^ row & wildtype – 2^nd^ row) and transgenic animals are shown for days 1, 4 and 7 for dorsal (left) and ventral (right). ‘s’ denotes the average of (number of synaptic puncta per 100 μm) from ‘n’ animals.

### Synaptotoxic effects are not specific to dysfunctional proteolysis of C99APP substrate

To determine if stalled E-S complex formed between mutant PSEN1 with a substrate other than C99 can affect proteolysis by γ-secretase, we created human Notch1-based transgenic DNA encoding a C-terminal fragment analogous to C99APP (C99Notch1) as well as its V50F/L51F double mutation, designed to block ɛ cleavage when either Phe is positioned at P2’ during proteolytic processing by the enzyme. We have recently reported using mass-spectrometry and ELISA that C99Notch1 V50F/L51F mutations result in ∼50% reduction in proteolysis of Notch-1 substrate by γ-secretase (Malvankar and Wolfe 2025), similar to but not as substantial as the near complete block of proteolysis of the analogous V50F/M51F in C99APP (Bolduc, Montagna et al. 2016). To test the effects of this double mutation on synaptic degeneration in *C. elegans*, expression vectors were designed to comprise of codons for signal peptide (as used in C99APP constructs) followed by 99 amino acids of the Notch1 extracellular truncation (NEXT) C-terminal fragment, generated from ligand-dependent juxtamembrane cleavage by ADAM-10 and a direct substrate for γ-secretase (Brou, Logeat et al. 2000, Mumm, Schroeter et al. 2000). wt C99Notch1 and mutant C99Notch1 transgenes were designed with *rgef-1* pan-neuronal promoter and *unc-54* 3’ UTR as used in C99APP expression vectors (see Supporting Information for full DNA sequences). Note that this truncated form of Notch1 does not contain regions responsible for nuclear localization and transcriptional regulation.

Co-expression of V50F/L51F C99Notch-1 with wt PSEN1 resulted in shorter median lifespan in both independent lines compared to *juIs1* and wt Notch-1 + wt PSEN1 (**Figure 7a & Figure S2a)**. The survival curves of V50F/M51F C99Notch-1 + wt PS-1 mutants versus wt C99Notch-1 + wt PS-1 were statistically different based on pairwise comparisons using Holm-Sidak method (**Table S1**). Synaptic loss occurred as early as day 1 in V50F/M51F C99Notch-1 animals co-expressing wt PSEN1 in both nerve cords (**Figure 7b-7c & Figure S2b-S2c)**. These results suggest that stalled γ-secretase complexes can be formed from dysfunctional proteolysis of a substrate other than C99APP and trigger synaptotoxicity in *C. elegans*.

**Figure 7.**
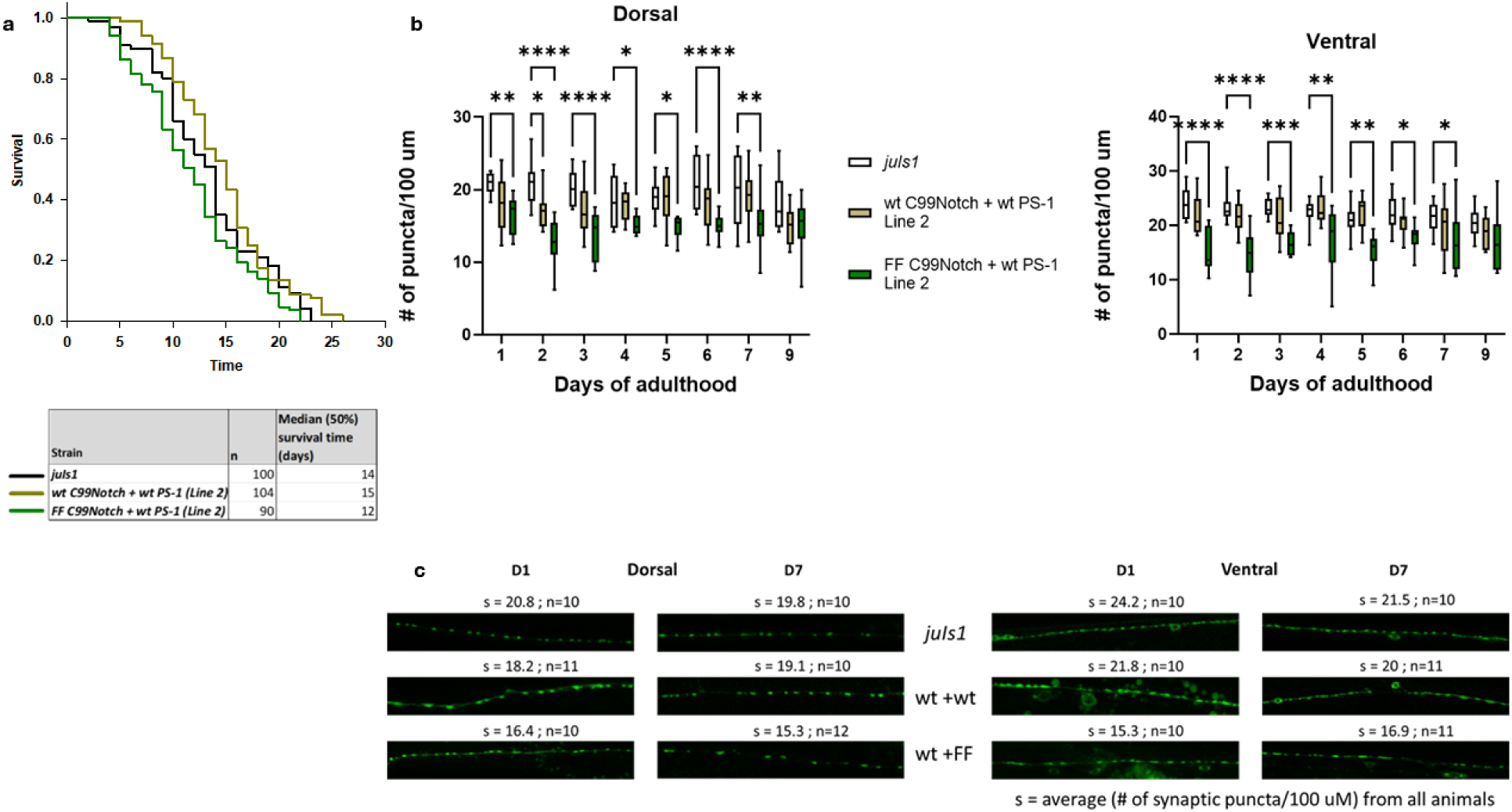
Expressing FFC99Notch with wt PS-1 triggers synaptic loss and shorter lifespan. **a. Lifespan of FF C99Notch + wt PS-1 animals** Kaplan-Meier curves of ‘n’ animals show the fraction of animals alive on different days. Age of control animals – juIs1 and wt C99Notch + wt PS-1 at median survival was 14 days while that of FF C99Notch + wt PS-1 was 13 days. The survival curve was statistically different between wt C99Notch + wt PS-1 and FF C99Notch + wt PS-1 animals (comparison table in supplementary). **b. Loss of synaptic puncta in FF C99Notch + wt PS-1 animals.** The average number of synaptic puncta per 100 μm in ‘n’ transgenic worms for each day is shown in the vertical box plots for dorsal and ventral cords. The horizontal lines in the top and bottom of any box plot represent the minimum and the values in that dataset of ‘n’ values. The ‘n’ for each day for a given mutant can be found in panel c. The upper and lower ends of a box mark the quartiles Q1 and Q3 values respectively. The horizontal line inside the box shows the median value (also, 2^nd^ quartile Q2). In comparison to *juIs1*, the number of synaptic puncta in FF C99Notch + wt PS-1 animals is lower from day 1 significantly until day 9in dorsal and day 7 in ventral nerve cords. (p>0.05 ns, p≤ 0.05 *, p≤ 0.01 **, p≤ 0.001 ***, p< 0.0001 ****). **c. Loss of synaptic puncta in FF C99Notch + wt PS-1 FAD animals.** 100 μm sections of representative confocal images for wildtype (1^st^ row) and transgenic animals are shown for days 1and 7 for dorsal (left) and ventral (right). ‘s’ denotes the average of (number of synaptic puncta per 100 μm) from ‘n’ animals. In FF C99Notch + wt PS-1 dorsal nerve cord, on day 1, s value is significantly lower (s=16.7), compared to s=20.8 in *juIs1*. Similarly, the value of s is significantly lower in mutant C99Notch (s=11.8) on day 7 compared to *juIs1*(s=19.8) in dorsal nerve. Similar representative images can be seen for ventral nerve cord. Note the gaps in synaptic puncta in these images, increasing the distance between two consecutive puncta.

### Catalytic aspartate mutation that eliminates proteolytic activity does not lead to reduced lifespan or synaptic degeneration

To test whether synaptic degeneration and reduction in lifespan is due to a simple loss of proteolytic function of γ-secretase, we utilized a PSEN1 transgene containing a mutation in one of the catalytic aspartates (D257A). Three independently derived lines containing the human D257A PS1 construct were obtained after microinjection and assessed for both synapse degeneration and lifespan. When assessed for median survival, none of the PSEN1 D257A mutant lines showed significantly reduced lifespan compared to parental strain *juIs1* (**Figure 8a; Table S1)**. WT PSEN1 + WT C99APP animals from strain *lhEx661* did have significantly longer median lifespans compared to parental line *juIs1* and two lines transgenic for D257A PS1. The third D257A PS1 line had a significantly longer lifespan compared to *juIs1* and to the other two lines containing the same transgene. This is likely due to variability in transgene expression between the three independently derived lines.

**Figure 8.**
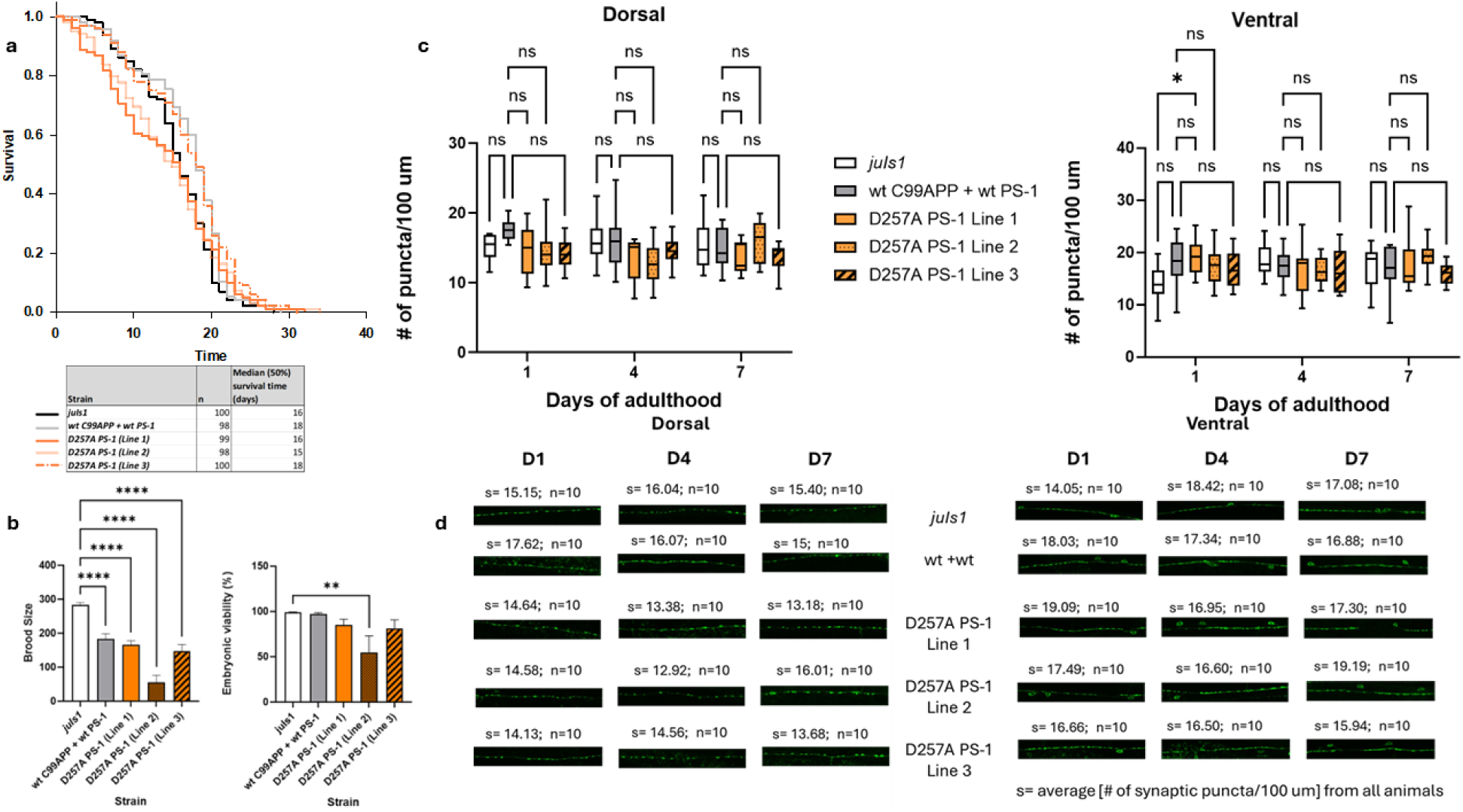
Catalytic aspartate mutations that eliminate proteolytic activity do not lead to a neurodegenerative phenotype. **a. Lifespan of D257A PS-1 animals** Kaplan-Meier curves of ‘n’ animals show the fraction of animals alive on different days. Age of control animals – *juIs1* and wt C99APP + wt PS-1 at median survival was 16 and 18 days respectively. The survival curve of D257A PS-1 Line 3 is statistically not different when compared to wt C99APP + wt PS-1 control but is statistically different when compared to survival curve of *juIs1* animals (comparison table in supplementary). **b. Brood size and embryonic viability of D257A PS-1 mutant animals. Final “n” values** for animals: juIs1 (10), WT PS1 + WT APP C99 (10), D257A 1 (9), D257A 2 (6), D257A 3 (10). Animals that died before the 5 days of scoring were excluded from final analysis. **c. Synaptic puncta quantification in D257A PS-1 animals.** The average number of synaptic puncta per 100 μm in ‘n’ transgenic worms for each day is shown in the vertical box plots for dorsal and ventral cords. The horizontal lines in the top and bottom of any box plot represent the minimum and the values in that dataset of ‘n’ values. The ‘n’ for each day for a given mutant can be found in panel d. The upper and lower ends of a box mark the quartiles Q1 and Q3 values respectively. The horizontal line inside the box shows the median value (also, 2^nd^ quartile Q2). In comparison to wt C99APP + wt PS-1 control, the number of synaptic puncta in D257A PS-1 animals is not significantly different on the days quantified (days 1, 4 & 7) in dorsal and ventral nerve cords. (p>0.05 ns, p≤ 0.05 *). **d. Synaptic puncta quantification in D257A PS-1 animals.** 100 μm sections of representative confocal images for controls (*juIs1* - 1^st^ row & wildtype – 2^nd^ row) and transgenic animals are shown for days 1, 4 and 7 for dorsal (left) and ventral (right). ‘s’ denotes the average of (number of synaptic puncta per 100 μm) from ‘n’ animals.

Interestingly, animals expressing D257A PSEN1 transgene had a slight visible egg-laying deficient phenotype. Egg-laying deficiency is present in animals with loss-of-function mutations in *sel-12*, a homolog of human Presenilin in *C. elegans*, and is linked to deficiencies in cleavage of NOTCH receptors (Levitan and Greenwald 1995). These visible defects prompted us to determine whether other aspects of nematode health, specifically brood size and embryonic viability, were affected. Animals expressing D257A PSEN1 showed significant differences in brood size, and one line showed significant defects in embryonic viability (**Figure 8b**). While the other two lines did not show significant changes in embryonic viability, they were slightly decreased compared to both *juIs1* and WT PSEN1 + WT APP C99 animals. In all lines, there were no overall significant differences between synapse numbers compared with parental line *juIs1* or wt C99APP + wt PSEN1 in either cord on any given day, indicating no age-dependent synapse degeneration (**Figure 8c**). Together, these results indicate that a simple loss of function of enzymatic activity is not sufficient to lead to synaptic degeneration and a reduced lifespan. The *sel-12*-like loss-of-function phenotypes (effects on brood size and embryonic viability) indicate that distinct pathways are involved.

## DISCUSSION

The amyloid cascade hypothesis of AD pathogenesis, first formulated over 30 years ago, implicates aggregated Aβ, particularly Aβ42, in triggering downstream neurotoxicity and synaptic dysfunction, ultimately leading to cognitive dysfunction. This hypothesis emerged with the discovery of FAD-associated dominant missense mutations in the APP gene, in and around the small Aβ region of the much larger precursor protein. The subsequent discovery of presenilin genes as sites of FAD mutations, that these mutations elevate the Aβ42/Aβ40 ratio, and that presenilins are proteases, the catalytic component of γ-secretase, seemingly cemented the amyloid hypothesis. All FAD mutations are found in the substrate and enzyme that produce Aβ, and these mutations alter the production or properties of Aβ peptides.

Decades later, fundamental gaps remain regarding the nature of pathogenic Aβ assemblies, the identity of cognate receptors, and pathways that relay synaptotoxic signals. In addition, monoclonal antibodies against Aβ aimed at halting the amyloid cascade and preventing neurodegeneration have shown only modest slowing of cognitive decline, despite effective clearance of amyloid plaques (Kurkinen, Fulek et al. 2023). For these reasons, we reinvestigated the identity of the pathogenic initiator in FAD. Through biochemical, structural, computational and cellular studies, we recently found that FAD mutations are deficient in multiple proteolytic steps in the processing of APP C99 substrate by γ-secretase due to stalling/stabilization of the enzyme-substrate (E-S) complex (Devkota, Zhou et al. 2024). Establishment of a *C. elegans* model system for FAD then provided a means of testing mechanisms of synaptotoxicity, leading to the conclusion that stalled γ-secretase E-S complexes, even in the absence of Aβ or in the presence of a substrate other than C99APP, can trigger synaptic loss.

In the present study, we provide important confirmation and extension of the “stalled complex” hypothesis of FAD pathogenesis (Wolfe 2025). Through the generation of independent transgenic *C. elegans* lines, we demonstrate that the FAD APP Iberian mutation I45F (Aβ numbering) elicits age-dependent synaptic loss and reduced lifespan that is largely dependent on co-expression of human wt PS1. This mutation leads to substantial elevation of the Aβ42/Aβ40 ratio, primarily by blocking γ-secretase trimming of Aβ43 to Aβ40 (Bolduc, Montagna et al. 2016). However, addition of the designed V44F mutation, which further blocks trimming of Aβ45 to Aβ42, does not rescue the neurodegenerative phenotype. Moreover, installation of the designed V50F/M51F double mutation near the initial γ-secretase cleavage site in APP C99—blocking all Aβ production even though this mutant substrate can nevertheless bind the protease complex—likewise leads to the neurodegenerative phenotype. These findings show that the age-dependent synaptic loss is not dependent on Aβ42 or overall Aβ production and are consistent with stalled E-S complexes as the pathogenic trigger.

We further observed that neuronal expression of FAD-mutant PSEN1 L166P leads to reduced lifespan in *C. elegans*, even in the absence of APP C99 coexpression, consistent with our previous report (Devkota, Zhou et al. 2024). In the present study, however, we observed the first disconnect in our new transgenic *C. elegans* FAD model system between reduced lifespan and age-dependent synaptic loss: Animals expressing PSEN1 L166P without wt C99APP expression showed reduced lifespan without synaptic degeneration. This new observation suggests that in this model system there may be multiple pathways and mechanisms by which PSEN1 FAD mutations might affect lifespan, many of which may not be coupled to synaptic degeneration. This finding further suggests that synaptic degeneration in this model system requires co-expression of endogenous substrate to increase levels of stalled E-S complexes.

We hypothesized that, with FAD-mutant PSEN1, formation of stalled γ-secretase E-S complexes with other substrates could trigger synaptic degeneration. Consistent with this idea, we observed in two independent transgenic lines that neuronal co-expression of WT PSEN1 and V50F/L51F Notch1-based substrate, which is cleaved 50% less efficiently by γ-secretase than its wt counterpart (Malvankar and Wolfe 2025), led to synaptic loss and a shorter median lifespan in our *in vivo* model. This shows that synaptic loss in *C. elegans* can be triggered by stalled E-S complexes with a substrate other than APP C99. This finding is consistent with a new study demonstrating that PSEN1 FAD mutations can cause neurodegeneration in the absence of APP in mice (Yan, Zhang et al. 2024). γ-Secretase has broad substrate specificity, with ∼150 known membrane protein substrates (Güner and Lichtenthaler 2020). We suggest that FAD-mutant PSEN1/γ-secretase can form stalled E-S complexes with all or most of these other substrates to elevate total levels of synaptotoxic stalled E-S complexes.

We further show in this study that a simple loss of γ-secretase catalytic activity is not sufficient to cause the observed synaptic loss and reduced lifespan seen with FAD mutants. Three transgenic *C. elegans* lines were generated that neuronally express PSEN1 D257A, which contains a mutation of one of the essential conserved aspartate residues in the active site, thereby rendering this mutant PSEN1 catalytically dead. This finding is consistent with two observations about human FAD. First, although over 300 FAD-associated mutations in PSEN1 have been identified, including many in and around the active site, neither of the two catalytic aspartates are mutated in any case of FAD. Indeed, no FAD mutation has been found that leads to complete loss of proteolytic activity. Second, dominant loss-of-function mutations in PSEN1 and other components of the γ-secretase complex, resulting in nonsense-mediated decay of the mRNA and haploinsufficiency, are associated with hereditary acne inverse, not neurodegeneration (Wang, Yang et al. 2010). Thus, although FAD mutations result in reduced proteolytic function, this partial loss of function alone apparently cannot elicit FAD. Presenilin proteolytic function does appear to be disrupted in the animals expressing PSEN1 D257A, as indicated by the observed *sel-12*-like egg-laying defect. Thus, loss of proteolytic function alone does not lead to age-dependent synaptic degeneration or reduce lifespan in this *C. elegans* model system. Whether co-expression of wt C99APP with PSEN1 D257A causes synaptic degeneration and reduces lifespan in these animals remains to be determined, and synergistic effects between reduction of proteolytic function and gain of new synaptotoxic function, both due to stalled γ-secretase E-S complexes, are still possible.

Our findings clearly point to stalled E-S complexes *per se* as an initiator of synaptic loss. In humans with AD, synaptic density correlates with cognitive function (DeKosky and Scheff 1990). The transgenic *C. elegans* system described here may provide a valuable model to understand mechanisms of synaptic dysfunction observed in all AD, not only FAD. In sporadic late-onset AD, with no mutation in APP or presenilin, other factors may lead to stalled γ-secretase E-S complexes, for instance changes in membrane composition or intracellular or intraorganelle pH. γ-Secretase is a membrane-embedded aspartyl protease that cleaves within the transmembrane domain of its many substrates. Altered cholesterol levels can affect γ-secretase activity in cells and proteoliposomes (Wahrle, Das et al. 2002, Osenkowski, Ye et al. 2008, Kim, Kim et al. 2016), and γ-secretase activity has long been known to be pH-dependent (Li, Lai et al. 2000, Fraering, Ye et al. 2004, Quintero-Monzon, Martin et al. 2011, Maesako, Houser et al. 2022). The effects of these factors on all γ-secretase proteolytic processing steps and E-S complex stability are currently under investigation. We are also searching for compounds that rescue stalled complexes, as a potential therapeutic strategy for the prevention or treatment of AD.

Given the advantages of *C. elegans* as a model for neurodegenerative studies, future research may leverage these FAD models for genetic screening to identify mediators of synaptic dysfunction. Studies in *C. elegans* continue to highlight conserved molecular pathways that contribute to synapse abnormalities and the relevance of this model organism for investigating pathological mechanisms of AD (Van Pelt and Truttmann 2020, Alvarez, Alvarez-Illera et al. 2022, Xu, Kang et al. 2025) and FAD (Sarasija, Laboy et al. 2018, Ashkavand, Ryan et al. 2025). Revealing molecular mechanisms of pathogenesis in *C. elegans* neurodegenerative models may then lead to development of therapeutic agents for the prevention and treatment of Alzheimer’s Disease. In this regard, the new *C. elegans* system described here may also provide a rapid and facile *in vivo* platform for AD drug discovery.

## METHODS

### *C. elegans* maintenance

All *C. elegans* strains used in the study were maintained at 15-22° C on nematode growth medium (NGM) plates seeded with *E. coli (Brenner 1974)*. *juIs1* [*Punc-25*::SNB-1::GFP] was used as the parental strain to develop transgenic lines by injecting human C99 APP and/or PSEN1 or human C99 Notch and/or PSEN1 constructs along with co-injection marker *rol-6* (Kramer, French et al. 1990).

### Cloning & molecular biology

Human C99APP and PS-1 cDNA were synthesized as described in a previous study (Devkota, Zhou et al. 2024). Similarly, human C99 Notch1 transgenes (wt and FF) were designed to include the following: 1) DNA encoding signal peptide (MLPGLALLLLAAWTARA) for membrane insertion, 2) coding sequence for 99 amino acids of Notch1 (wt and FF). These constructs were designed to include worm introns. All human constructs were cloned in worm expression vector pEVL415 (P*rgef-1*:*htau40::gfp::unc-54* 3′UTR) (Devkota, Zhou et al. 2024). Restriction digestion was performed using BamH1 and NgoM4 to linearize the vector. Plasmids harboring C99 wt Notch1 and C99 FF Notch1 sequences analogous to C99 APP constructs used in the previous study were synthesized using ‘GeneArt Gene Synthesis’ by ThermoFisher. These plasmids contain Kan^R^ genes for growth on bacterial plates and DNA extraction. The genes of interest were PCR amplified from the Thermofisher plasmids using primers that were designed to have ∼ 15 bp 5’ extensions complementary to the 3’ sticky ends of the linearized vector. The fragments were PCR-purified and cloning was performed by In-Fusion cloning (TakaraBio) following manufacturer’s protocol. Ampicillin resistance allowed for specific isolation of bacterial clones containing Notch constructs. Clones were confirmed by PCR amplification and restriction digestion. In addition, plasmid DNA containing the gene inserts were validated by sequencing.

### *C. elegans* transgenesis

The parental strain *juIs1* [*unc-25*p::*snb-1*::GFP + lin-15(+)] IV expresses GFP fused to synaptobrevin in presynaptic terminals of GABAergic DD and VD motor neurons and RME neurons (Hallam and Jin 1998). Human constructs with co-injection marker were microinjected into gonads of young adults of *juIs1* [*Punc-25*::SNB-1::GFP] to obtain transgenic lines. Injection mixes contain human APP (or Notch1) variant (8-10 ng/μl) and/ or human PS-1 variant (8-10 ng/μl) and pRF4 (60.8 ng/μl). Injection mixes for D257A PS1 contained 10ng/µl of DNA and pRF4 (60 ng/µl). Plasmids used in the study are below. Full sequences of new transgenes are provided as a supplementary files **Text S1**, **Text S2** and **Text S3**).

1. pEVL545 [*Prgef-1*:signal peptide : human wt C99APP :: *unc-54* 3’UTR]
2. pEVL546 [*Prgef-1*::signal peptide : human C99APP (I45F) :: *unc-5*4 3’UTR
3. pEVL547 [*Prgef-1*::signal peptide : human wt PS-1 :: *unc-54* 3’UTR]
4. pEVL548 [*Prgef-1*:: human L166P PS-1 :: *unc-54* 3’UTR]
5. pEVL549 [*Prgef-1*::signal peptide : human V44F I45F C99APP :: *unc-54* 3’UTR]
6. pEVL554 [*Prgef-1*::signal peptide : human V50F M51F C99APP :: *unc-54* 3’UTR]
7. pEVL555 [*Prgef-1*::signal peptide : human wt C99Notch :: *unc-54* 3’UTR] (**Text S1**)
8. pEVL556 [*Prgef-1*::signal peptide : human FF C99 Notch :: *unc-54* 3’UTR] (**Text S2**)
9. pEVL553 [P*rgef-1*:: human D257A PS-1 :: *unc-54* 3’UTR] (**Text S3**)

Strains generated in this study

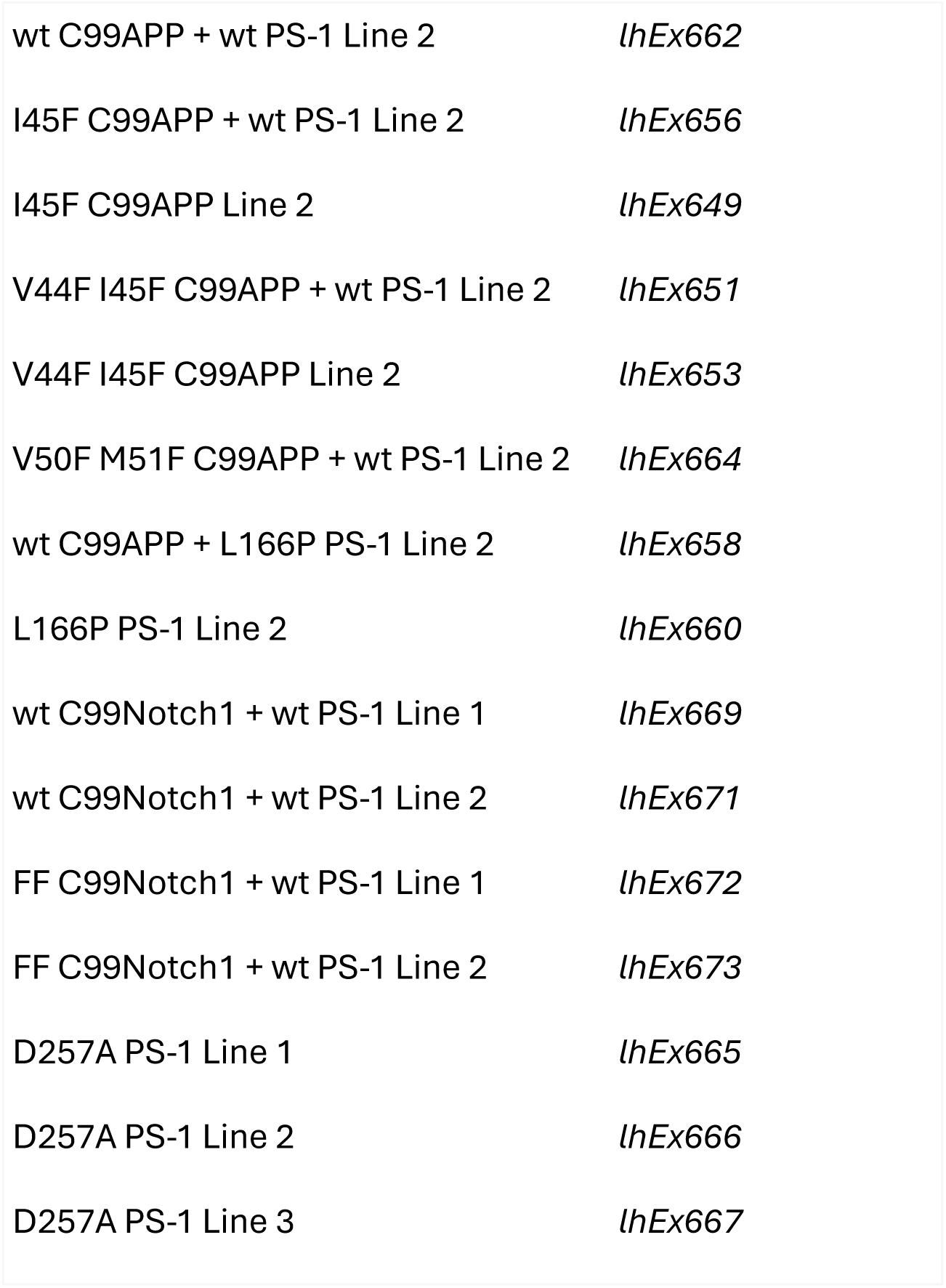

### Lifespan

L4 animals were isolated for lifespan experiments and maintained at 20 °C. To avoid contamination with progeny animals, adults were transferred to fresh plates whenever necessary. Plates were checked for live worms at least once in two days. Animals that did not move when prodded were considered dead.

### Microscopy and image analysis

L4 animals were transferred to fresh plates and were maintained at 20 °C. Day 1 to day 7 adults and day 9 adults were scored for the number of SNB-1::GFP puncta in the ventral and dorsal nerve cords. 0.5% 2-phenoxypropanol in M9 was used to anesthetize adult worms and those were mounted on 2% agarose pads for imaging. Imaging was done using Olympus FV1000 laser scanning confocal microscope at 60X magnification NA 1.42. All images were obtained processed and analyzed as described previously (Devkota, Zhou et al. 2024).

*C. elegans* were also imaged in the Microscopy and Analytical Imaging Resource Core Laboratory (RRID:SCR_021801) at The University of Kansas using a TCS SPE Laser Scanning Confocal Upright Microscope (Leica Microsystems, DM6-Q model), with the 488 nm laser line, a Leica 63X/1.3NA ACS APO oil objective, 12-bit spectral PMT detector and a Leica LAS X Imaging software (version 3.5.7.23225). Synaptic puncta signal was detected using 488 nm excitation, 500-520 nm emission range. Images were captured at 1024 x 1024-pixel resolution, no bidirectional scanning, and a zoom factor at 1.0.

### Agarose Immobilization

For imaging of animals containing the D257A PS1 and L166P PS1 transgene, immobilization with agarose and polystyrene beads was used. L4 animals were transferred to fresh plates and maintained at 20°C. 10% agarose in M9 buffer was used to make the agarose pads. 1.5 uL of 0.1 um polystyrene beads (Sigma-Aldrich catalog 90517, Batch BCCL4220) were added to immobilize worms. Day 1, day 4, and day 7 adults were scored for the number of SNB-1::GFP puncta in the ventral and dorsal nerve cords in this manner. All control animals were also scored using this method and compared against D257A and L166P mutants. Image instrumentation and analysis was the same as all other lines imaged under anesthetization using the Olympus FV1000.

### Lifespan for D257A

L4 animals were isolated and maintained at 20 °C for lifespan experiments. Animals were picked and plated in technical replicates of 4 plates equaling a total number of 100 animals at minimum. Animals were transferred to new plates and scored for life every day to avoid contamination. Animals that did not move when prodded were considered dead.

### Brood size & Embryonic Viability Assays

To analyze the brood size, on day 1, one L4 hermaphrodite was transferred to each plate. The worms were allowed to develop into adults and lay eggs for 24 hours at 20 °C, after which the adult worm was transferred to a new plate on day 2. It was ensured that no eggs or other worms were transferred inadvertently. On day 3, the day 1 plate was scored separately for L1 larvae and unhatched eggs. This process was repeated until day 5 when the adult worm was removed and day 7 when the last scoring was recorded for day 5 plates (Hornsten, Lieberthal et al. 2007, Gallrein, Iburg et al. 2021).

### Data analysis

#### Lifespan

Plotting of survival curves, calculation of median lifespan and statistical analysis were performed in ‘Sigmaplot’ using Kaplan Meier (log-rank test) method.

#### Microscopic image analysis

All confocal image processing and analysis were performed using Fiji (ImageJ) software, as described in our previous report (Devkota, Zhou et al. 2024).

#### Brood size & embryonic viability

Brood size for each replicate of a given strain is the sum of total progeny including unhatched embryos (along with live larvae) from each parent hermaphrodite, obtained by summing the daily counts from day 1 to day 5.

Brood size = Live progeny + unhatched embryo.

Embryonic viability percentage was calculated for each biological replicate of a given strain The numbers of live progeny and unhatched embryos are obtained by totaling the daily count across the experimental period (Kwah and Jaramillo-Lambert 2023).

Embryonic viability = [Total live progeny/ (Total live progeny + Total unhatched embryo)] × 100

In both cases, statistical analysis was performed in GraphPad Prism using one way ANOVA test (Dunnett’s multiple comparisons test). Comparisons were made with *juIs1* with p>0.05 ns, p≤ 0.05 *, p≤ 0.01 **, p≤ 0.001 ***, p< 0.0001 ****

## Acknowledgment

We thank E. Lundquist (U. Kansas) for injection apparatus for microinjection of transgenes into *C. elegans*, and E. Rosa-Molinar and N. Martinez-Rivera for confocal microscopy images obtained at the Microscopy and Analytical Imaging Research Resource Core Laboratory at U. Kansas. This work was supported by grants AG66986 and AG79569 from the US National Institutes of Health and a Pilot Project Grant from the University of Kansas Alzheimer Disease Research Center via NIH grant P30 AG072973.

## REFERENCES

Adamla, F. and Z. Ignatova (2015). “Somatic expression of unc-54 and vha-6 mRNAs declines but not pan-neuronal rgef-1 and unc-119 expression in aging Caenorhabditis elegans.” Sci Rep 5: 10692.

Alvarez, J., P. Alvarez-Illera, J. Santo-Domingo, R. I. Fonteriz and M. Montero (2022). “Modeling Alzheimer’s Disease in Caenorhabditis elegans.” Biomedicines 10(2).

Arafi, P., S. Devkota, E. Williams, M. Maesako and M. S. Wolfe (2025). “Alzheimer-mutant γ-secretase complexes stall amyloid β-peptide production.” Elife 13: RP102274.

Ashkavand, Z., K. C. Ryan, J. T. Laboy, R. Patel, B. Geller and K. R. Norman (2025). “Identification of presenilin mutations that have sufficient gamma-secretase proteolytic activity to mediate Notch signaling but disrupt organelle and neuronal health.” Neurobiol Dis 212: 106961.

Bateman, R. J., P. S. Aisen, B. De Strooper, N. C. Fox, C. A. Lemere, J. M. Ringman, S. Salloway, R. A. Sperling, M. Windisch and C. Xiong (2011). “Autosomal-dominant Alzheimer’s disease: a review and proposal for the prevention of Alzheimer’s disease.” Alzheimers Res Ther 3(1): 1.

Bieschke, J., E. Cohen, A. Murray, A. Dillin and J. W. Kelly (2009). “A kinetic assessment of the C. elegans amyloid disaggregation activity enables uncoupling of disassembly and proteolysis.” Protein Sci 18(11): 2231–2241.

Bolduc, D. M., D. R. Montagna, M. C. Seghers, M. S. Wolfe and D. J. Selkoe (2016). “The amyloid-beta forming tripeptide cleavage mechanism of gamma-secretase.” Elife 5.

Breijyeh, Z. and R. Karaman (2020). “Comprehensive Review on Alzheimer’s Disease: Causes and Treatment.” Molecules 25(24).

Brenner, S. (1974). “The genetics of Caenorhabditis elegans.” Genetics 77(1): 71–94.

Brou, C., F. Logeat, N. Gupta, C. Bessia, O. LeBail, J. R. Doedens, A. Cumano, P. Roux, R. A. Black and A. Israël (2000). “A Novel Proteolytic Cleavage Involved in Notch Signaling: The Role of the Disintegrin-Metalloprotease TACE.” Molecular Cell 5: 207–216.

Cai, Q. and P. Tammineni (2017). “Mitochondrial Aspects of Synaptic Dysfunction in Alzheimer’s Disease.” J Alzheimers Dis 57(4): 1087–1103.

Chen, L., Y. Fu, M. Ren, B. Xiao and C. S. Rubin (2011). “A RasGRP, C. elegans RGEF-1b, couples external stimuli to behavior by activating LET-60 (Ras) in sensory neurons.” Neuron 70(1): 51–65.

DeKosky, S. T. and S. W. Scheff (1990). “Synapse loss in frontal cortex biopsies in Alzheimer’s disease: correlation with cognitive severity.” Ann Neurol 27(5): 457–464.

Devkota, S., M. Maesako and M. S. Wolfe (2025). “Presenilin-1 Familial Alzheimer Mutations Impair γ-Secretase Cleavage of APP Through Stabilized Enzyme–Substrate Complex Formation.” Biomolecules 15: 955.

Devkota, S., T. D. Williams and M. S. Wolfe (2021). “Familial Alzheimer’s disease mutations in amyloid protein precursor alter proteolysis by γ-secretase to increase amyloid β-peptides of >45 residues.” J Biol Chem. 296:100281.(doi): 10.1016/j.jbc.2021.100281.

Devkota, S., R. Zhou, V. Nagarajan, M. Maesako, H. Do, A. Noorani, C. Overmeyer, S. Bhattarai, J. T. Douglas, A. Saraf, Y. Miao, B. D. Ackley, Y. Shi and M. S. Wolfe (2024). “Familial Alzheimer mutations stabilize synaptotoxic gamma-secretase-substrate complexes.” Cell Rep 43(2): 113761.

Fraering, P. C., W. Ye, J. M. Strub, G. Dolios, M. J. LaVoie, B. L. Ostaszewski, A. Van Dorsselaer, R. Wang, D. J. Selkoe and M. S. Wolfe (2004). “Purification and Characterization of the Human gamma-Secretase Complex.” Biochemistry 43(30): 9774–9789.

Gallrein, C., M. Iburg, T. Michelberger, A. Kocak, D. Puchkov, F. Liu, S. M. Ayala Mariscal, T. Nayak, G. S. Kaminski Schierle and J. Kirstein (2021). “Novel amyloid-beta pathology C. elegans model reveals distinct neurons as seeds of pathogenicity.” Prog Neurobiol 198: 101907.

Goate, A., M. C. Chartier-Harlin, M. Mullan, J. Brown, F. Crawford, L. Fidani, L. Giuffra, A. Haynes, N. Irving, L. James and et al. (1991). “Segregation of a missense mutation in the amyloid precursor protein gene with familial Alzheimer’s disease.” Nature 349(6311): 704–706.

Güner, G. and S. F. Lichtenthaler (2020). “The substrate repertoire of γ-secretase/presenilin.” Semin Cell Dev Biol 105: 27–42.

Hallam, S. J. and Y. Jin (1998). “lin-14 regulates the timing of synaptic remodelling in Caenorhabditis elegans.” Nature 395(6697): 78–82.

Hornsten, A., J. Lieberthal, S. Fadia, R. Malins, L. Ha, X. Xu, I. Daigle, M. Markowitz, G. O’Connor, R. Plasterk and C. Li (2007). “APL-1, a Caenorhabditis elegans protein related to the human beta-amyloid precursor protein, is essential for viability.” Proc Natl Acad Sci U S A 104(6): 1971–1976.

Hunter, P. (2024). “The controversy around anti-amyloid antibodies for treating Alzheimer’s disease : The European Medical Agency’s ruling against the latest anti-amyloid drugs highlights the ongoing debate about their safety and efficacy.” EMBO Rep.

Kim, Y., C. Kim, H. Y. Jang and I. Mook-Jung (2016). “Inhibition of Cholesterol Biosynthesis Reduces γ-Secretase Activity and Amyloid-β Generation.” J Alzheimers Dis 51(4): 1057–1068.

Kramer, J. M., R. P. French, E. C. Park and J. J. Johnson (1990). “The Caenorhabditis elegans rol-6 gene, which interacts with the sqt-1 collagen gene to determine organismal morphology, encodes a collagen.” Mol Cell Biol 10(5): 2081–2089.

Krstic, D. and I. Knuesel (2013). “Deciphering the mechanism underlying late-onset Alzheimer disease.” Nat Rev Neurol 9(1): 25–34.

Kurkinen, M., M. Fulek, K. Fulek, J. A. Beszlej, D. Kurpas and J. Leszek (2023). “The Amyloid Cascade Hypothesis in Alzheimer’s Disease: Should We Change Our Thinking?” Biomolecules 13(3).

Kwah, J. K. and A. Jaramillo-Lambert (2023). “Measuring Embryonic Viability and Brood Size in Caenorhabditis elegans.” J Vis Exp(192).

Levitan, D., T. G. Doyle, D. Brousseau, M. K. Lee, G. Thinakaran, H. H. Slunt, S. S. Sisodia and I. Greenwald (1996). “Assessment of normal and mutant human presenilin function in Caenorhabditis elegans.” Proc Natl Acad Sci U S A 93(25): 14940–14944.

Levitan, D. and I. Greenwald (1995). “Facilitation of lin-12-mediated signalling by sel-12, a Caenorhabditis elegans S182 Alzheimer’s disease gene.” Nature 377(6547): 351–354.

Li, Y. M., M. T. Lai, M. Xu, Q. Huang, J. DiMuzio-Mower, M. K. Sardana, X. P. Shi, K. C. Yin, J. A. Shafer and S. J. Gardell (2000). “Presenilin 1 is linked with gamma-secretase activity in the detergent solubilized state.” Proc Natl Acad Sci U S A 97(11): 6138–6143.

Link, C. D. (1995). “Expression of human beta-amyloid peptide in transgenic Caenorhabditis elegans.” Proc Natl Acad Sci U S A 92(20): 9368–9372.

Maesako, M., M. C. Q. Houser, Y. Turchyna, M. A.-O. Wolfe and O. Berezovska (2022). “Presenilin/γ-Secretase Activity Is Located in Acidic Compartments of Live Neurons.” J Neurosci 42(1): 145–154.

Makin, S. (2018). “The amyloid hypothesis on trial.” Nature 559(7715): S4–S7.

Malvankar, S. R. and M. S. Wolfe (2025). “Effects of Transmembrane Phenylalanine Residues on gamma-Secretase-Mediated Notch-1 Proteolysis.” ACS Chem Neurosci 16(5): 844–855.

McColl, G., B. R. Roberts, A. P. Gunn, K. A. Perez, D. J. Tew, C. L. Masters, K. J. Barnham, R. A. Cherny and A. I. Bush (2009). “The Caenorhabditis elegans A beta 1-42 model of Alzheimer disease predominantly expresses A beta 3-42.” J Biol Chem 284(34): 22697–22702.

McColl, G., B. R. Roberts, T. L. Pukala, V. B. Kenche, C. M. Roberts, C. D. Link, T. M. Ryan, C. L. Masters, K. J. Barnham, A. I. Bush and R. A. Cherny (2012). “Utility of an improved model of amyloid-beta (Abeta(1)(-)(4)(2)) toxicity in Caenorhabditis elegans for drug screening for Alzheimer’s disease.” Mol Neurodegener 7: 57.

Merritt, C., D. Rasoloson, D. Ko and G. Seydoux (2008). “3’ UTRs are the primary regulators of gene expression in the C. elegans germline.” Curr Biol 18(19): 1476–1482.

Morris, J. C., M. Weiner, C. Xiong, L. Beckett, D. Coble, N. Saito, P. S. Aisen, R. Allegri, T. L. S. Benzinger, S. B. Berman, N. J. Cairns, M. C. Carrillo, H. C. Chui, J. P. Chhatwal, C. Cruchaga, A. M. Fagan, M. Farlow, N. C. Fox, B. Ghetti, A. M. Goate, B. A. Gordon, N. Graff-Radford, G. S. Day, J. Hassenstab, T. Ikeuchi, C. R. Jack, W. J. Jagust, M. Jucker, J. Levin, P. Massoumzadeh, C. L. Masters, R. Martins, E. McDade, H. Mori, J. M. Noble, R. C. Petersen, J. M. Ringman, S. Salloway, A. J. Saykin, P. R. Schofield, L. M. Shaw, A. W. Toga, J. Q. Trojanowski, J. Vöglein, S. Weninger, R. J. Bateman and V. D. Buckles (2022). “Autosomal dominant and sporadic late onset Alzheimer’s disease share a common in vivo pathophysiology.” Brain 145(10): 3594–3607.

Mumm, J. S., E. H. Schroeter, M. T. Saxena, A. Griesemer, X. Tian, D. J. Pan, W. J. Ray and R. Kopan (2000). “A Ligand-Induced Extracellular Cleavage Regulates Gamma-Secretase-like Proteolytic Activation of Notch1.” Molecular Cell 5: 197–206.

Munoz-Jimenez, C., C. Ayuso, A. Dobrzynska, A. Torres-Mendez, P. C. Ruiz and P. Askjaer (2017). “An Efficient FLP-Based Toolkit for Spatiotemporal Control of Gene Expression in Caenorhabditis elegans.” Genetics 206(4): 1763–1778.

Osenkowski, P., W. Ye, R. Wang, M. S. Wolfe and D. J. Selkoe (2008). “Direct and potent regulation of gamma-secretase by its lipid microenvironment.” J Biol Chem 283(33): 22529–22540.

Pelucchi, S., F. Gardoni, M. Di Luca and E. Marcello (2022). “Synaptic dysfunction in early phases of Alzheimer’s Disease.” Handb Clin Neurol 184: 417–438.

Pope, C. A., H. M. Wilkins, R. H. Swerdlow and M. S. Wolfe (2021). “Mutations in the Amyloid-beta Protein Precursor Reduce Mitochondrial Function and Alter Gene Expression Independent of 42-Residue Amyloid-beta Peptide.” J Alzheimers Dis 83(3): 1039–1049.

Quintero-Monzon, O., M. M. Martin, M. A. Fernandez, C. A. Cappello, A. J. Krzysiak, P. Osenkowski and M. S. Wolfe (2011). “Dissociation between the processivity and total activity of gamma-secretase: implications for the mechanism of Alzheimer’s disease-causing presenilin mutations.” Biochemistry 50(42): 9023–9035.

Sarasija, S., J. T. Laboy, Z. Ashkavand, J. Bonner, Y. Tang and K. R. Norman (2018). “Presenilin mutations deregulate mitochondrial Ca(2+) homeostasis and metabolic activity causing neurodegeneration in Caenorhabditis elegans.” Elife 7: e33052.

Sherrington, R., E. I. Rogaev, Y. Liang, E. A. Rogaeva, G. Levesque, M. Ikeda, H. Chi, C. Lin, G. Li, K. Holman, T. Tsuda, L. Mar, J. F. Foncin, A. C. Bruni, M. P. Montesi, S. Sorbi, I. Rainero, L. Pinessi, L. Nee, I. Chumakov, D. Pollen, A. Brookes, P. Sanseau, R. J. Polinsky, W. Wasco, H. A. Da Silva, J. L. Haines, M. A. Perkicak-Vance, R. E. Tanzi, A. D. Roses, P. E. Fraser, J. M. Rommens and P. H. St George-Hyslop (1995). “Cloning of a gene bearing missense mutations in early-onset familial Alzheimer’s disease.” Nature 375(6534): 754–760.

Subramanian, J., J. C. Savage and M. E. Tremblay (2020). “Synaptic Loss in Alzheimer’s Disease: Mechanistic Insights Provided by Two-Photon in vivo Imaging of Transgenic Mouse Models.” Front Cell Neurosci 14: 592607.

Takami, M., Y. Nagashima, Y. Sano, S. Ishihara, M. Morishima-Kawashima, S. Funamoto and Y. Ihara (2009). “gamma-Secretase: successive tripeptide and tetrapeptide release from the transmembrane domain of beta-carboxyl terminal fragment.” J Neurosci 29(41): 13042–13052.

Tanzi, R. E. (2012). “The genetics of Alzheimer disease.” Cold Spring Harb Perspect Med 2(10): pii: a006296.

Treusch, S., S. Hamamichi, J. L. Goodman, K. E. Matlack, C. Y. Chung, V. Baru, J. M. Shulman, A. Parrado, B. J. Bevis, J. S. Valastyan, H. Han, M. Lindhagen-Persson, E. M. Reiman, D. A. Evans, D. A. Bennett, A. Olofsson, P. L. DeJager, R. E. Tanzi, K. A. Caldwell, G. A. Caldwell and S. Lindquist (2011). “Functional links between Abeta toxicity, endocytic trafficking, and Alzheimer’s disease risk factors in yeast.” Science 334(6060): 1241–1245.

Van Pelt, K. M. and M. C. Truttmann (2020). “Caenorhabditis elegans as a model system for studying aging-associated neurodegenerative diseases.” Transl Med Aging 4: 60–72.

Wahrle, S., P. Das, A. C. Nyborg, C. McLendon, M. Shoji, T. Kawarabayashi, L. H. Younkin, S. G. Younkin and T. E. Golde (2002). “Cholesterol-dependent gamma-secretase activity in buoyant cholesterol-rich membrane microdomains.” Neurobiol Dis 9(1): 11–23.

Wang, B., W. Yang, W. Wen, J. Sun, B. Su, B. Liu, D. Ma, D. Lv, Y. Wen, T. Qu, M. Chen, M. Sun, Y. Shen and X. Zhang (2010). “Gamma-secretase gene mutations in familial acne inversa.” Science 330(6007): 1065.

Wolfe, M. S. (2025). “Presenilin, γ-Secretase, and the Search for Pathogenic Triggers of Alzheimer’s Disease.” Biochemistry 64(8): 1662–1672.

Xu, R., Q. Kang, X. Yang, P. Yi and R. Zhang (2025). “Unraveling Molecular Targets for Neurodegenerative Diseases Through Caenorhabditis elegans Models.” Int J Mol Sci 26(7).

Yan, K., C. Zhang, J. Kang, P. Montenegro and J. Shen (2024). “Cortical neurodegeneration caused by Psen1 mutations is independent of Abeta.” Proc Natl Acad Sci U S A 121(34): e2409343121.

Zhang, H., Q. Ma, Y. W. Zhang and H. Xu (2012). “Proteolytic processing of Alzheimer’s beta-amyloid precursor protein.” J Neurochem 120 Suppl 1(Suppl 1): 9–21.

