## Supplementary material for "A *C. elegans* model of familial Alzheimer’s disease shows age-dependent synaptic degeneration independent of amyloid β-peptide": Figure S1, Figure S2, Figure S3, Table S1, Text S1, Text S2, Text S3

**Supplemental Materials**

1. **Supplemental Figures**

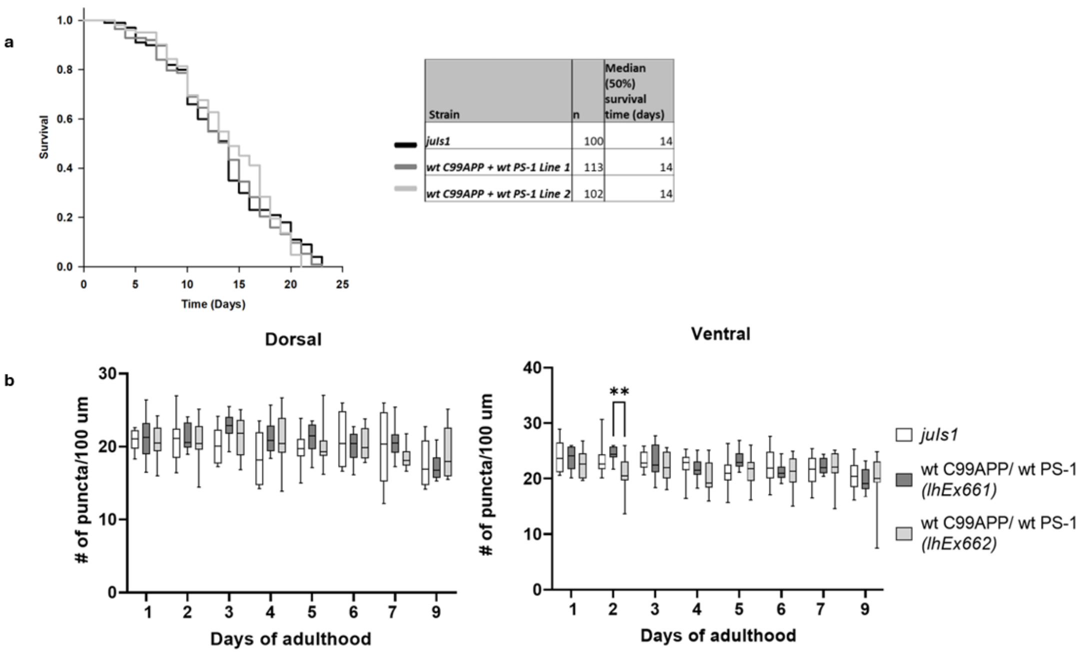

| Strain | n | Median (50%) survival time (days) |
|---|---|---|
| *juls1* | 100 | 14 |
| *wt C99APP + wt PS-1 Line 1* | 113 | 14 |
| *wt C99APP + wt PS-1 Line 2* | 102 | 14 |

**Figure S1**

a. Lifespan of *juls1* and wt C99APP + wt PS-1 animals. Kaplan-Meier curves of 'n' animals show the fraction of animals alive on different days. The median age of parent line *juls1* animals was not significantly different from that of any of the wt C99APP + wt PS-1 lines.

b. Synaptic puncta of *juls1* and wt C99APP + wt PS-1 animals. The average number of synaptic puncta per 100 μm in 'n' transgenic worms for each day is shown in the vertical box plots for dorsal and ventral cords. The horizontal lines in the top and bottom of any box plot represent the minimum and the values in that dataset of 'n' values. The 'n' for each day for a given mutant can be found in figure d. The upper and lower ends of a box mark the quartiles Q1 and Q3 values respectively. The horizontal line inside the box shows the median value (also, 2nd quartile Q2). The number of synaptic puncta in dorsal and ventral nerve cords of both wt C99APP + wt PS-1 lines were comparable to their parent line *juls1* on all days studied. ($p>0.05$ ns, $p \leq 0.05$ *, $p \leq 0.01$ **, $p \leq 0.001$ ***, $p < 0.0001$ ****)

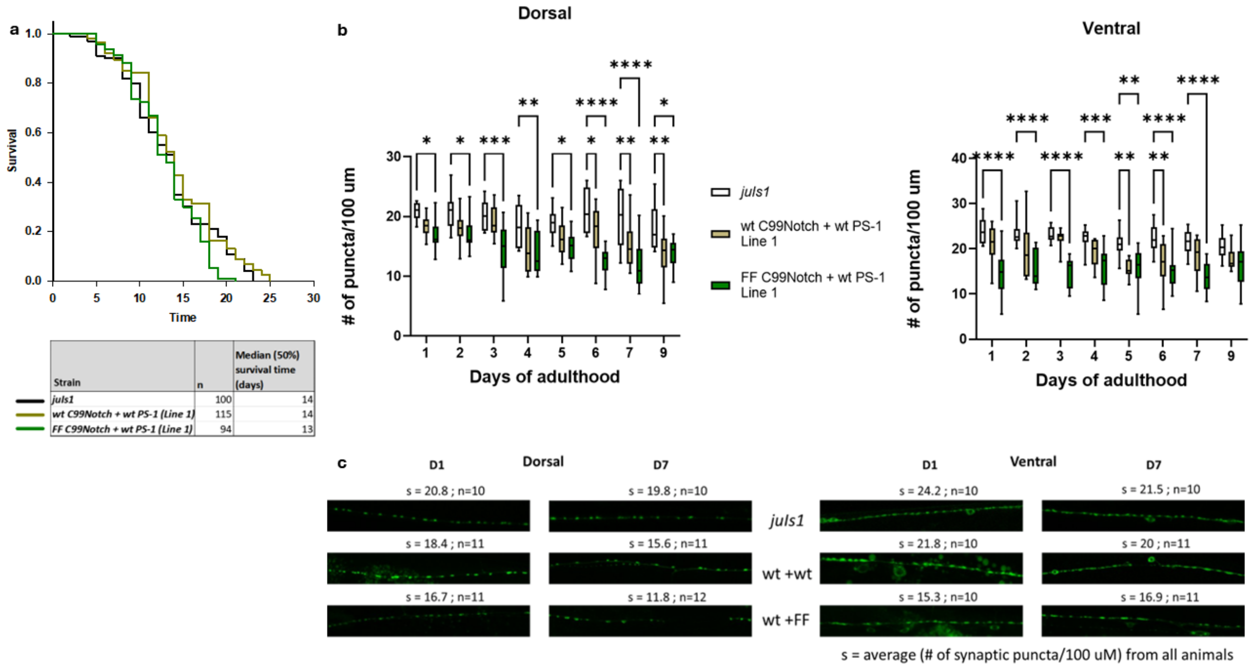

**Figure S2**

a. Lifespan of FF C99Notch + wt PS-1 (Line 1) animals. Kaplan-Meier curves of 'n' animals show the fraction of animals alive on different days. Age of control animals – *juls1* and wt C99Notch + wt PS-1 at median survival was 14 and 15 days respectively while that of FF C99Notch + wt PS-1 was 12 days. The survival curves were statistically different between control and FF C99Notch + wt PS-1 animals.

b. Loss of synaptic puncta in FF C99Notch + wt PS-1 animals. The average number of synaptic puncta per 100 μm in 'n' transgenic worms for each day is shown in the vertical box plots for dorsal and ventral cords. The horizontal lines in the top and bottom of any box plot represent the minimum and the values in that dataset of 'n' values. The 'n' for each day for a given mutant can be found in figure f. The upper and lower ends of a box mark the quartiles Q1 and Q3 values respectively. The horizontal line inside the box shows the median value (also, 2$^{nd}$ quartile Q2). In comparison to *juls1*, the number of synaptic puncta in FF C99Notch + wt PS-1 animals is lower from day 1 significantly until day 7 in dorsal and ventral nerve cords. ($p>0.05$ ns, $p\leq 0.05$ *, $p\leq 0.01$ **, $p\leq 0.001$ ***, $p< 0.0001$ ****).

c. Loss of synaptic puncta in FF C99Notch + wt PS-1 FAD animals. 100 μm sections of representative confocal images for wildtype (1$^{st}$ row) and transgenic animals are shown for days 1and 7 for dorsal (left) and ventral (right). 's' denotes the average of (number of synaptic puncta per 100 μm) from 'n' animals. In FF C99Notch + wt PS-1 dorsal nerve cord, on day 1, s value is significantly lower (s=16.4), compared to s=20.8 in *juls1*. Similarly, the value of s is significantly lower in mutant C99Notch (s=15.3) on day 7 compared to *juls1*(s=19.8) in dorsal nerve. Similar representative images can be seen for ventral nerve cord. Note the gaps in synaptic puncta in these images, increasing the distance between two consecutive puncta.

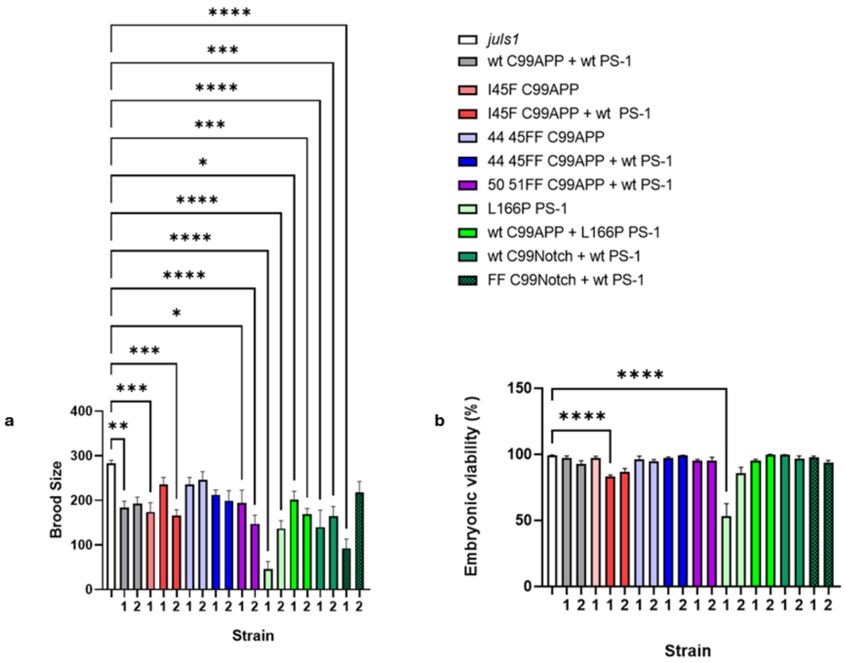

**Figure S3**

a. The column bars represent mean brood size values ± SEM.
b. The column bars represent mean embryonic viability values ± SEM.

### 2. Supplemental Table

| ***Table S1:*** *Pairwise comparisons of survival curves of C. elegans transgenic lines. All pairwise multiple comparisons used the Holm-Sidak method. Overall significance level cutoff <0.05* | | |
|---|---|---|
| Pairwise comparison of survival curves of *C. elegans* transgenic lines using data from experiments shown in main Figure 2. | | |
| **Comparisons** | **P Value** | **Significant?** |
| wt C99APP + wt PS-1 vs I45F C99APP + wt PS-1 | 1.109E-13 | Yes |
| wt C99APP + wt PS-1 vs. I45F C99APP | 0.0000272 | Yes |
| I45F C99APP + wt PS-1 vs. I45F C99APP | 0.0000161 | Yes |
| Pairwise comparison of survival curves of *C. elegans* transgenic lines using data from experiments shown in main Figure 3. | | |
| **Comparisons** | **P Value** | **Significant?** |
| wt C99APP + wt PS-1 vs. V44F I45F C99APP + wt PS-1 | 4.53E-08 | Yes |
| wt C99APP + wt PS-1 vs. V44F I45F C99APP | 0.239 | No |
| V44F I45F C99APP + wt PS-1 vs. V44F I45F C99APP | 0.0000491 | Yes |
| Pairwise comparison of survival curves of *C. elegans* transgenic lines using data from experiments shown in main Figure 4. | | |
| **Comparisons** | **P Value** | **Significant?** |
| wt C99APP + wt PS-1 vs. V50F M51F C99APP + wt PS-1 | 0.00897 | Yes |
| I45F C99APP + wt PS-1 vs. V50F M51F C99APP + wt PS-1 | 0.0176 | Yes |
| V44F I45F C99APP + wt PS-1 vs. V50F M51F C99APP + wt PS-1 | 0.481 | No |
| Pairwise comparison of survival curves of *C. elegans* transgenic lines using data from experiments shown in main Figure 5. | | |
| **Comparisons** | **P Value** | **Significant?** |
| wt C99APP + wt PS-1 vs. wt C99APP + L166P PS-1 | 7.261E-14 | Yes |
| wt C99APP + wt PS-1 vs. L166P PS-1 | 2.3E-09 | Yes |
| wt C99APP + L166P PS-1 vs. L166P PS-1 | 0.0381 | Yes |
| Pairwise comparison of survival curves of *C. elegans* transgenic lines using data from experiments shown in main Figure 7. | | |
| **Comparisons** | **P Value** | **Significant?** |
| *juIs1* vs. [wt C99Notch + wt PS-1 (Line 2)] | 0.0709 | No |
| *juIs1* vs. [FF C99Notch + wt PS-1 (Line 2)] | 0.0441 | Yes |
| [wt C99Notch + wt PS-1 (Line 2)] vs. [FF C99Notch + wt PS-1 (Line 2)] | 0.00133 | Yes |

| Pairwise comparisons of survival curves of *C. elegans* transgenic lines using data from experiments shown in Figure S1. | | |
|---|---|---|
| **Comparisons** | **P Value** | **Significant?** |
| *juIs1* vs. [wt C99Notch + wt PS-1 (Line 1)] | 0.213 | No |
| *juIs1* vs. [FF C99Notch + wt PS-1 (Line 1)] | 0.123 | No |
| [wt C99Notch + wt PS-1 (Line 1)] vs. [FF C99Notch + wt PS-1 (Line 1)] | 0.0186 | Yes |
| Pairwise comparison of survival curves of *C. elegans* transgenic lines using data from experiments shown in main Figure 8. | | |
| **Comparisons** | **P Value** | **Significant?** |
| *juIs1* vs. wt C99APP + wt PS-1 | 0.0773 | No |
| *juIs1* vs. D257A Line 1 | 0.935 | No |
| *juIs1* vs. D257A Line 2 | 0.992 | No |
| *juIs1* vs. D257A Line 3 | 0.046 | Yes |
| wt C99APP + wt PS-1 Line 1 vs. D257A Line 1 | 0.469 | No |
| wt C99APP + wt PS-1 Line 1 vs. D257A Line 2 | 0.487 | No |
| wt C99APP + wt PS-1 Line 1 vs. D257 Line 3 | 0.84 | No |
| D257A Line 1 vs. D257A Line 2 | 0.994 | No |
| D257A Line 1 vs. D257A Line 3 | 0.122 | No |
| D257A Line 2 vs. D257A Line 3 | 0.225 | No |

### 3. Supplemental *C. elegans* transgene sequences

Note: Underlined– codons for the signal peptide ; Lower case– C. elegans introns

**Text S1.** pEVL555 [P*rgef-1*::signal peptide : human wt C99 Notch-1 :: *unc-54* 3'UTR]

ATGCTGCCCGGTTTGGCACTGCTCCTGCTGGCCGCCTGGACGGCTCGGGCGGTGCAGA
GTGAGACCGTGGAGCCGCCCCCGCCGGCGCAGCTGCACTTCATgtaagtttaaacatatatata
ctaactaaccctgattatttaaattttcagGTACGTGGCGGCGGCCGCCTTTGTGCTTCTGTTCTTCGT
GGGCTGCGGGGTGCTGCTGTCCCGCAAGCGCCGGCGGCAGCATGGCCAGCTCTGGTT
CCCTGAGGGCTTCAAAGTgtaagtttaaacatatatatactaactaaccctgattatttaaattttcagGTCTGA
GGCCAGCAAGAAGAAGCGGCGGGAGCCCCTCGGCGAGGACTCCGTGGGCCTCAAGC
CCCTGAAGAACGCTTCAGACGGTGCCCTCATGGACGACAACCAGAATGAGTGGGGGGAC
GAGGACCTGGAGTAG

**Text S2.** pEVL556 [P*rgef-1*::signal peptide : human FFC99 Notch-1 :: *unc-54* 3'UTR]

ATGCTGCCCGGTTTGGCACTGCTCCTGCTGGCCGCCTGGACGGCTCGGGCGGTGCAGA
GTGAGACCGTGGAGCCGCCCCCGCCGGCGCAGCTGCACTTCATgtaagtttaaacatatatata

ctaactaaccctgattatttaaattttcagGTACGTGGCGGCGGCCGCCTTTGTGCTTCTGTTCTTCGT
GGGCTGCGGGTTCTTTCTGTCCCGCAAGCGCCGGCGGCAGCATGGCCAGCTCTGGTTC
CCTGAGGGCTTCAAAGTgtaagtttaaacatatatatactaactaaccctgattatttaaattttcagGTCTGAG
GCCAGCAAGAAGAAGCGGCGGGAGCCCCTCGGCGAGGACTCCGTGGGCCTCAAGCC
CCTGAAGAACGCTTCAGACGGTGCCCTCATGGACGACAACCAGAATGAGTGGGGGGACG
AGGACCTGGAGTAG

**Text S3.** pEVL553 [P*rgef-1*:: human D257A PS-1 :: *unc-54* 3'UTR]

ATGACAGAGTTACCTGCACCGTTGTCCTACTTCCAGAATGCACAGATGTCTGAGGACAACC
ACCTGAGCAATACTGTACGTAGCCAGAATGACAATAGAGAACGGCAGGAGCACAACGACA
GACGGAGCCTTGGCCACCCTGAGCCATTATCTAATGGACGACCCCAGGGTAACTCCCGG
CAGGTGGTGGAGCAAGATGAGGAAGAAGATGAGGAGCTGACATTGAAATATGGCGCCAAG
CATGTGATCATGCTCTTTGTCCCTGTGACTCTCTGCATGGTGGTGGTCGTGGCTACCATTAAG
gtaagtttaaacatatatatactaactaaccctgattatttaaattttcagTCAGTCAGCTTTTATACCCGGAAG
GATGGGCAGCTAATCTATACCCCATTCACAGAAGATACCGAGACTGTGGGCCAGAGAGCC
CTGCACTCAATTCTGAATGCTGCCATCATGATCAGTGTCATTGTTGTCATGACTATCCTCCTG
GTGGTTCTGTATAAATACAGGTGCTATAAGGTCATCCATGCCTGGCTTATTATATCATCTCTATT
GTTGCTGTTCTTTTTTTCATTCATTTACTTGGGGGAAGTGTTTAAAACCTATAACGTTGCTGTGG
ACTACATTACTGTTGCACTCCTGATCTGGAATTTTGGTGTGGTGGGAATGATTTCCATTCACTG
GAAAGGTCCACTTCGACTCCAGCAGgtaagtttaaacagttcggtactaactaaccatacatatttaaatttt
cagGCATATCTCATTATGATTAGTGCCCTCATGGCCCTGGTGTTTATCAAGTACCTCCCTGAAT
GGACTGCGTGGCTCATCTTGGCTGTGATTTCAGTATATGCATTAGTGGCTGTTTTGTGTCCGA
AAGGTCCACTTCGTATGCTGGTTGAAACAGCTCAGGAGAGAAATGAAACGCTTTTTCCAGCT
CTCATTTACTCCTCAACAATGGTGTGGTTGGTGAATATGGCAGAAGGAGACCCGGAAGCTC
AAAGGAGAGTATCCAAAAATTCCAAGTATAATGCAGAAAGCACAGAAAGGGAGTCACAAGA
CACTGTTGCAGAGAATGATGATGGCGGGTTCAGTGAGGAATGGGAAGCCCAGAGGGACA
GTCATCTAGGGCCTCATCGCTCTACACCTGAGTCACGAGCTGCTGTCCAGGAACTTTCCAG
CAGTATCCTCGCTGGTGAAGACCCAGAGGAAAGGGGAGTAAAACTTGGATTGGGAGATTTC
ATTTTCTACAGTGTTCTGGTTGGTAAAGCCTCAGCAACAGCCAGTGGAGACTGGgtaagtttaa
acagttcggtactaactaaccatacatatttaaattttcagAACACAACCATAGCCTGTTTCGTAGCCATAT
TAATTGGTTTGTGCCTTACATTATTACTCCTTGCCATTTTCAAGAAAGCATTGCCAGCTCTTCC
AATCTCCATCACCTTTGGGCTTGTTTTCTACTTTGCCACAGATTATCTTGTACAGCCTTTTATGG
ACCAATTAGCATTCCATCAATTTTATATCTAG
